# Generation, characterization and drug sensitivities of twelve patient-derived IDH1 mutant glioma cell cultures

**DOI:** 10.1101/2021.04.09.435131

**Authors:** Cassandra Verheul, Ioannis Ntafoulis, Trisha V. Kers, Youri Hoogstrate, Pier G. Mastroberardino, Sander Barnhoorn, César Payán-Gómez, Romain Tching Chi Yen, Eduard A. Struys, Stijn L.W. Koolen, Clemens M.F. Dirven, Sieger Leenstra, Pim J. French, Martine L.M. Lamfers

**Author notes:** Both authors contributed equally to this manuscript (Pim J. French, Martine L.M. Lamfers). Correspondence, Wytemaweg 80, Ee2236, 3015 CN Rotterdam. **Authorship** Conceptualization: C.V., M.L.M.L. and P.J.F.; Methodology: C.V.; M.L.M.L., P.J.F.; Software: Y.H.; R.T.C.Y.; Formal analysis: Y.H., C.V., P.J.F., R.T.C.Y., C.P.G.; Investigation: C.V., I.N., T.V.K., S.B., E.A.S.; Resources: P.G.M., E.A.S.; Writing – Original Draft: C.V., I.N., M.L.M.L.; Writing – Review and editing: M.L.M., P.J.F., P.G.M., C.P.G., C.M.F.D., S.L.W.K., S.L.; Visualization: C.V.; Supervision: M.L.M.L., P.J.F., S.L.; Funding acquisition: M.L.M.L. and P.J.F.

## Abstract

**Background:** Mutations of the isocitrate dehydrogenase (*IDH*) gene occur in over 80% of low-grade gliomas and secondary glioblastomas. Despite considerable efforts, endogenous *in vitro IDH*-mutated glioma models remain scarce. Availability of these models is key for the development of new therapeutic interventions.

**Methods:** Cell cultures were established from fresh tumor material and expanded in serum-free culture media. D-2-Hydroxyglutarate levels were determined by mass-spectrometry. Genomic and transcriptomic profiling were carried out on the Illumina Novaseq platform, methylation profiling was performed with the Infinium MethylationEpic BeadChip array. Mitochondrial respiration was measured with the Seahorse XF24 Analyzer. Drug screens were performed with an NIH FDA-approved anti-cancer drug set and two IDH-mutant specific inhibitors.

**Results:** A set of twelve patient-derived IDHmt cell cultures was established. We confirmed high concordance in driver mutations, copy number and methylation profiles between the tumors and derived cultures. Homozygous deletion of *CDKN2A/B* was observed in all cultures. IDH-mutant cultures had lower mitochondrial reserve capacity. IDH-mutant specific inhibitors did not affect cell viability or global gene expression. Screening of 107 FDA-approved anti-cancer drugs identified nine compounds with potent activity against IDHmt gliomas, including three compounds with favorable pharmacokinetic characteristics for CNS penetration: teniposide, omacetaxine mepesuccinate, and marizomib.

**Conclusions:** Our twelve IDH-mutant cell cultures show high similarity to the parental tissues and offer a unique tool to study the biology and drug sensitivities of high-grade IDHmt gliomas *in vitro*. Our drug screening studies reveal lack of sensitivity to IDHmt inhibitors, but sensitivity to a set of nine available anti-cancer agents.

**Key points:**

1. IDHmt glioma cultures closely resemble their parental tumors
2. Microscopic monitoring of early passages and colony isolation increases IDH1mt culture success
3. Drug screening identified nine candidate repurposed drugs for IDHmt glioma

**Importance of the study:** IDH-mutations are highly prevalent in low grade and secondary high-grade gliomas. Despite this high frequency however, very few *in vitro* models have been reported for IDH-mutated gliomas. In this manuscript we describe and characterize in detail twelve primary cultures from IDH-mutant astrocytomas. We show that these cultures retain most of the genetic, epigenetic and metabolic features of their respective parental tumors. Because of these similarities, these independent model systems will not only help understand the molecular defects driven by the mutation, but are also vital to identify means to target these tumors. Screening of 107 FDA-approved anti-cancer agents on these cultures identified a set of highly effective agents that may offer candidates for either systemic or assisted delivery treatment of this tumor subtype.

## Background

*IDH* mutations are present in approximately 80% of low-grade glioma (grade II and III) and secondary glioblastomas.^1^ *IDH* mutations result in a neomorphic gain-of-function of the mutant enzyme, which causes it to convert alpha ketoglutarate (α-KG) to D-2-hydroxyglutarate (D-2-HG). D-2-HG is a competitive inhibitor of several α-KG dependent enzymes including the histone lysine demethylase JMJD2 and the methylcytosine dioxygenase TET2, which results in the global hypermethylated (G-CIMP, glioma CpG island methylator phenotype) state of IDH-mutant gliomas.^2,3^

The discovery of this somatic mutation in malignant glioma instigated the development of agents targeting *IDH*-mutant tumors. ^4,5^ Unfortunately, pre-clinical research has been hampered by lack of IDHmt glioma model systems, as tumor samples from IDH-mutated glioma patients are notoriously difficult to culture.^6,7^ Worldwide, only a few endogenous IDH-mutant cell lines have been described thus far, some derived from fresh patient material, others from patient-derived xenografts that were serially transplanted in mice.^8–14^

Over the last ten years, our lab has attempted to establish cell cultures from over 275 low-grade and secondary gliomas resected in our clinic. Our continuous effort resulted in 12 cultures from IDH-mutant 1p19q non-codeleted astrocytomas, which is the largest set of patient-derived IDHmt cell cultures described to date. We show that these cell cultures retain the morphological, genetic, epigenetic, metabolic and transcriptomic features of the primary tumor. Current developments in the neuro-oncology field directed toward improving drug delivery to CNS tumors have reignited interest in anti-cancer agents that are already available. We therefore applied our IDHmt cell cultures to screen 107 FDA-approved anti-cancer agents to identify potential candidates for either systemic or enhanced delivery applications.

## Methods

### Tumor processing and cell culture

Fresh glioma tissue samples were obtained directly from the operating room of the Erasmus Medical Center or the Elizabeth Tweesteden Hospital, the Netherlands. The use of patient tissue for this study was approved by the local ethics committees of these hospitals and all patients signed informed consent forms according to the guidelines of the Institutional Review Boards of the respective hospitals.

Samples were processed essentially according to a further optimized protocol based on Balvers et al (see also supplemental methods).^6^ After 5-8 days, cultures were transferred to a new flask coated with 1:100 Cultrex PathClear RGF-BME (R&D Systems) for adherent expansion. Cell cultures were considered successful if they could be passaged at least five times while retaining their *IDH1* mutation.

Growth rates and doubling times were assessed by seeding 2*10^5^ cells in a pre-coated T75 flasks (+/− 10 μM AGI-5198, Agios), which were counted and split at ~90% confluency for at least three passages.

### Image analysis of low passage cell populations

Cells were plated in a 96 wells CellCarrier Ultra plates (Perkin Elmer, Hamburg, Germany). After 6 days of culturing, live cells were stained with Hoechst and cell tracker green (1:1000, Thermo Fisher, Bleiswijk, the Netherlands) and imaged using a spinning disk Opera Phenix confocal high throughput microscope system (Perkin Elmer, Hamburg, Germany) equipped with a water immersion 40x objective. Images were analyzed using Harmony Analysis Software (PerkinElmer, Hamburg, Germany).

### Sequencing

Genomic DNA was extracted from cell pellets (passages ranging from 0-12), cryosections of snap frozen tumor material or leukocytes using the DNeasy Blood & Tissue Kit (Qiagen).

All RNA was extracted from pellets of cultured cells (passage numbers ranging from 8-14) and cryosections of snap frozen tumor material with the RNeasy Plus Mini Kit (Qiagen).

Presence of IDH1 mutations was confirmed by Sanger sequencing using 5’-GTG GCA CGG TCT TCA GAG A-3’ and 5’-TTC ATA CCT TGC TTA ATG GGT GT-3’ primers.

Five hundred ng genomic DNA was fragmented to ~300 bp with the Kapa Hyperplus Library prep kit. The exome captured by the SeqCap wash kit and SeqCap pure capture bead kit (both from Roche). The Illumina Novaseq platform was used to sequence 150 bases (paired-end sequencing) to obtain 6GB per sample.

Driver mutations were derived from.^15^ CopyNumbers were calculated using the CNVKit package.^16^ RNA was isolated from cell cultures (+/− 5 μM AGI-5198) using the RNeasy kit (Qiagen). We performed paired-end sequencing of 2×100 with the Illumina Novaseq platform to obtain 8-10 GB per sample. For details on sequencing, data processing and analysis, see supplemental methods.

DNA methylation levels was assessed using the Infinium methylationEPIC beadchip arrays (Illumina) according to standard protocols and classified as described ^17^ or using the TCGAbiolinks Bioconductor package. Further analysis as done using the Bioconductor *Minfi* package.^18^ The sequencing and gene expression data is available at Zenodo.org (doi: 10.5281/zenodo.4498024).

### Mitochondrial oxidative respiration assay

We measured cellular oxidative respiration with the Seahorse XF24 Analyzer as described previously.^19^ Cell cultures were seeded in ten-plo at a density of 40,000 cells per well and grown overnight in standard culture medium. One hour before the start of the experiment we replaced culture medium with XF Assay Medium (Agilent Technologies) pH7.4, supplemented with 10 mM glucose, 2 mM glutamin and 1 mM sodium pyruvate and incubated at 37°C without CO_2_. After baseline measurements, cellular response after sequential injections of 1 μM oligomycin, 0.5 μM FCCP, and 1 μM antimycin were measured. Basal respiration, mitochondrial ATP production, proton leakage, and maximal respiration rates were calculated with Seahorse Wave software.

### D/L-2-hydroxyglutarate measurements

Intracellular D-2-HG and L-2-HG was measured in cell pellets after five days of culture as described.^20^

### Drug screening

Drug screening was done using IDH inhibitors AGI-5198 (Agios) or BAY-1436032 (Bayer) or the FDA-approved Oncology Drug Set II library (National Cancer Institute). Viability was assessed by the CellTiter GLO 2.0. For details see supplemental methods.

### Gene expression analysis

Differential gene expression analysis and normalization was performed with DESeq2 and further visualizations of expression profiles used the DESeq2 VST transformed read counts ^21^. only genes with ≥4 read counts across the tested samples were included.

Principal component analysis was performed on the top 500 genes with the highest variance. We performed cluster analysis based on the 1000 most variably expressed genes using the pheatmap package in R, and scaled by row.

Pathway analysis was done using the DAVID Bioinformatics Resources 6.8 (Huang DW, Sherman BT, Lempicki RA. Systematic and integrative analysis of large gene lists using DAVID Bioinformatics Resources. Nature Protoc. 2009;4(1):44-57) using a false discovery rate (FDR) of 0.05 as cut-off and using Gene Set Enrichment Analysis (GSEA) using an FDR < 0.25.^22^ Statistical significance of pathway enrichment scores was ascertained by permutation testing with matched random gene sets.

### Statistical analysis

The unpaired Student’s T-test was used for the comparison of two groups (statistical significance was defined as p<0.05). To determine significance of contingency tables we used the Fisher exact test (statistical significance was defined as P<0.05). The IC50 values were calculated by applying a Nonlinear regression (curve-fit) and selecting the dose-response inhibition equation in Graphpad Prism.

## Results

### Establishment of successful IDHmt cell cultures

Glioma resection material of over 275 low-grade or secondary high-grade gliomas were transported to our lab to establish patient-derived cell cultures. We sequenced all successful cultures (*N*=80) for IDH mutations and identified twelve cell cultures derived from eleven patients that retained their IDH-R132H mutation. Nine cultures were derived from secondary glioblastomas, one from a grade III anaplastic astrocytoma and two from grade II astrocytomas. All cultures were derived from 1p/19q non-codeleted gliomas. Patient and cell line characteristics are listed in Table S1. Table S2 summarizes on which levels each IDHmt cell culture has been characterized.

The doubling time of IDHmt cultures (mean 5.2 days; range 1.8-16.4) was longer than of IDHwt glioblastoma cultures (mean 2.6 days; range 1.6-3.6, p<0.05), albeit not significant (Fig. S1A). We passaged GS.0580 to passage 50 without any notable changes in growth rate or morphology and verified the continued presence of the *IDH1* mutation. We determined intracellular levels of the enantiomers D-2-HG and L-2-HG in all our IDHmt and seven IDHwt cell cultures. All IDHmt cell cultures revealed dramatically increased levels of D-2-HG over L-2-HG (mean value 859; range 21-2700) compared to those of IDHwt cell cultures (mean value 1.2; range 0.3-5.3, p<0.05), indicating presence of mutant IDH enzyme activity (Fig. S1B).

### IDHmt cultures resemble parental tumors on genomic level

We performed whole exome sequencing on seven IDHmt cultures and their parental tumors to determine whether they retained the classical features of IDHmt astrocytic gliomas. Apart from the *IDH1* mutation, *TP53* was most frequently mutated in the cell cultures (N=7), followed by *ATRX* (Fig. 1A). Both driver mutations were retained in cell cultures if they were present in parental tumors.

**Figure 1.**
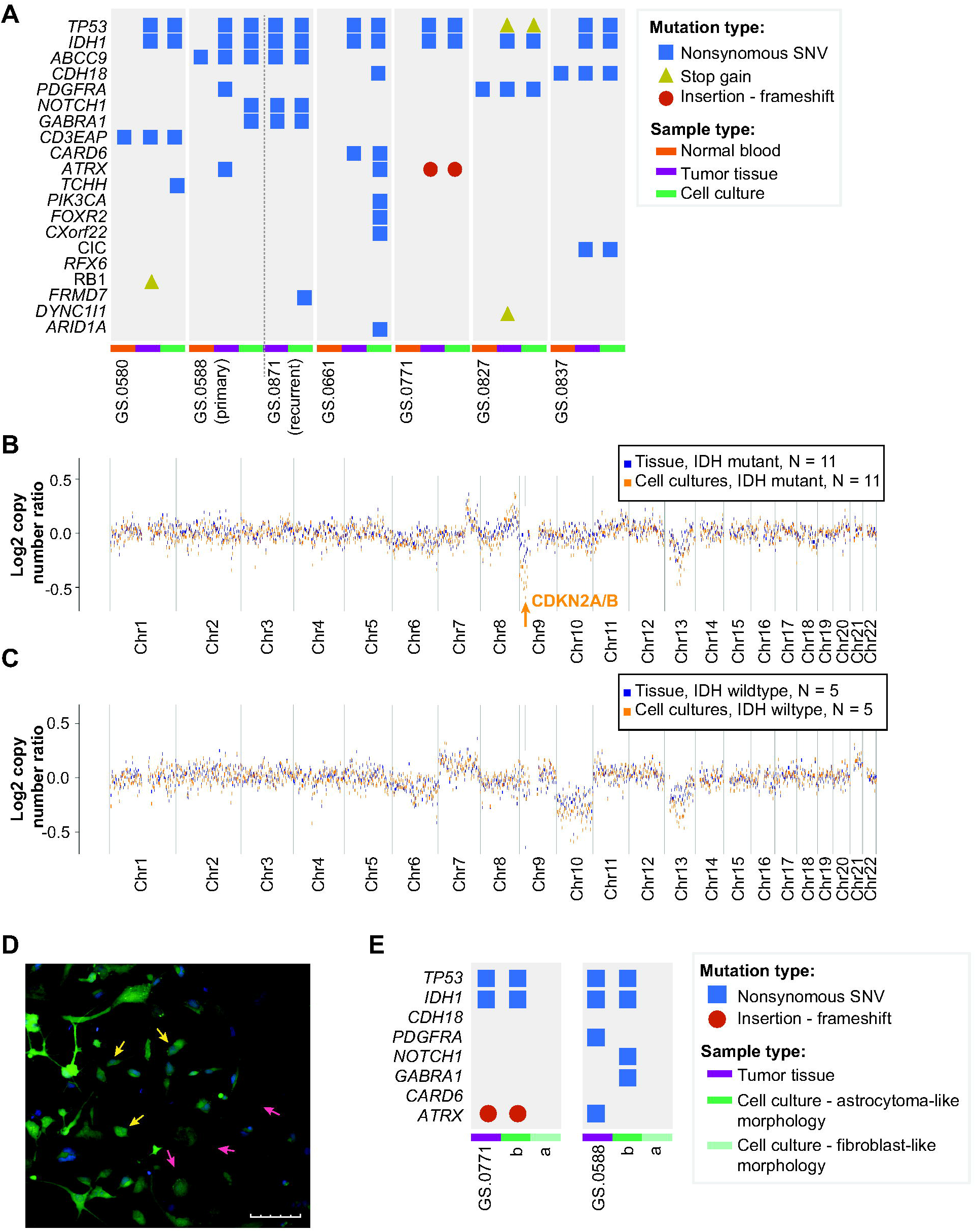
IDHmt cell cultures represent the genotype of IDHmt gliomas. **(A)** Presence of glioma-associated mutations in non-neoplastic control tissue (orange bars), tumor tissue (purple bars), or cell cultures (green bars). Mutation types included are nonsynonymous SNVs (blue squares), stop gains (yellow triangles) and frameshift (red circles). **(B, C)** Median copy number log ratio plots of IDHmt tissues and cultures (N=11), and IDHwt tissues and cultures (N=5) based on global methylation data. On the X-axes chromosomes, on the Y-axes median log ratio of copy number change. Chromosomal gains and losses are more pronounced in cultures than in tissues, probably due to the presence of non-cancerous cells in tumor tissue. **(D)** Fluorescent image of an early IDH mutant cell culture with two distinct phenotypes: astrocytoma-like (three individual cells pointed out by yellow arrows) and fibroblast-like (three individual cells pointed out by pink arrows). Scale bar represents 100 micron. **(E)** Identified mutations in IDHmt tumors GS.0771 and GS0588 and derived daughter cultures, one with astrocytoma-like and one with fibroblast-like morphology.

Interestingly, mutations in *NOTCH1* and *GABRA1* were not identified in one primary tumor sample, while they were found in the primary cell culture (GS.0588b). At tumor recurrence, however, both mutations were present in the tumor and cell culture, which indicates subclonal expansion of mutations already detectable in our primary cell culture. Furthermore, in copy number analysis we found a striking homozygous loss of the *CDKN2A/B* locus in all IDHmt cell cultures, whereas this loss was detected in only eight out of twelve IDHmt tumors (Fig. 1B, Fig. S2A-L). When compared to the glioblastomas present in the TCGA database, *CDKN2A/B* loss in combination with IDH1/2 mutations was significantly more common in our cell cultures (12/12 compared to 6/15, p=0.001). We did not note such specific copy number changes when we compared IDHwt glioma cultures to their parental tumors.

Overall, copy number analysis shows similarity of copy number alterations between the tumor tissues and corresponding cell cultures for both IDHmt and IDHwt cultures (Fig. S2).

### 2D culture morphology reveals IDH status

In early passages of the IDHmt cell cultures often two types of cells were visible: round-bodied cells with thin protrusions and bright edges (astrocytic phenotype) and dark, large flat cells (fibroblast-like phenotype). We hypothesized that IDHmt glioma cells would be of the astrocytic phenotype. We separated morphologically different colonies from low-passage (≤1) cell cultures. Colony isolation and expansion from a mixed population of cells, yielded two distinct cell cultures: the original culture (designated ‘a’, e.g., GS.0771a) that evolved to almost exclusively contain fibroblast-like cells, and an astrocytoma-like culture (designated ‘b’). The two distinct cell types can be observed in a mix culture in Fig. 1D). Sanger sequencing revealed that only GS.0771b maintained its IDH mutation and copy numbers of the parental tumor; whereas GS.0771a did not retain the IDH mutation and showed a diploid genome (Fig. S3 A-C). Mutations in *TP53* and *ATRX* from the parental tumor were preserved in GS.0771b but not GS.0771a (Fig. 1E). Similarly, clone separation of GS.0588, also showed retention of *IDH1* and *TP53* mutations in the astrocytic GS.0588b culture but not in the fibroblast-like GS0588a culture. All successful cultures with verified IDH mutations had astrocytoma-like morphologies, although some degree of heterogeneity between cultures was observed (Fig. S4A-L). They are readily distinguishable from the fibroblast-like cultures that no longer harbor the IDH mutation by brightfield microscopy (Fig S4M-O). Quantification of morphologic parameters of the fibroblast-like and astrocytoma-like phenotype are summarized in table S3. The most distinguishing factor is cell area (6.1·10^3^ μm^2^ for IDHwt vs. 1.2·10^3^ μm^2^ for IDHmt), which is the main feature that can be identified by eye. Furthermore, the nuclei of IDHmt glioma cultures are significantly smaller, and the GFP intensity is higher when live-staining the cells (Fig 1D). Thus, astrocytoma-like morphological features can be used as pre-selection for establishing IDHmt cell cultures.

### Global methylation profiles are retained in IDHmt glioma cultures

To determine whether the epigenetic profiles of our cultures were retained, we assessed the DNA methylation class of twelve IDHmt and 5 IDHwt cell cultures and their parental tumors. In a publicly available classifier that can predict all CNS tumor types, all cell cultures were classified as gliomas and received the same overall class as their parental tumors with a classifier score above the cutoff of 0.9, except for glioma cell culture GS.0811 which received a score of 0.74.^17^ All IDHmt tumors were classified as IDHmt with subclass either A_IDH (astrocytoma, IDH mutant, N=4) or A_IDH_HG (astrocytoma, IDHmt high grade, N=8) (Fig. 2A). Interestingly, all IDHmt cultures were assigned to methylation subclass A_IDH_HG which points towards tumor progression in culture or to selection of more aggressive subclones.

**Figure 2.**
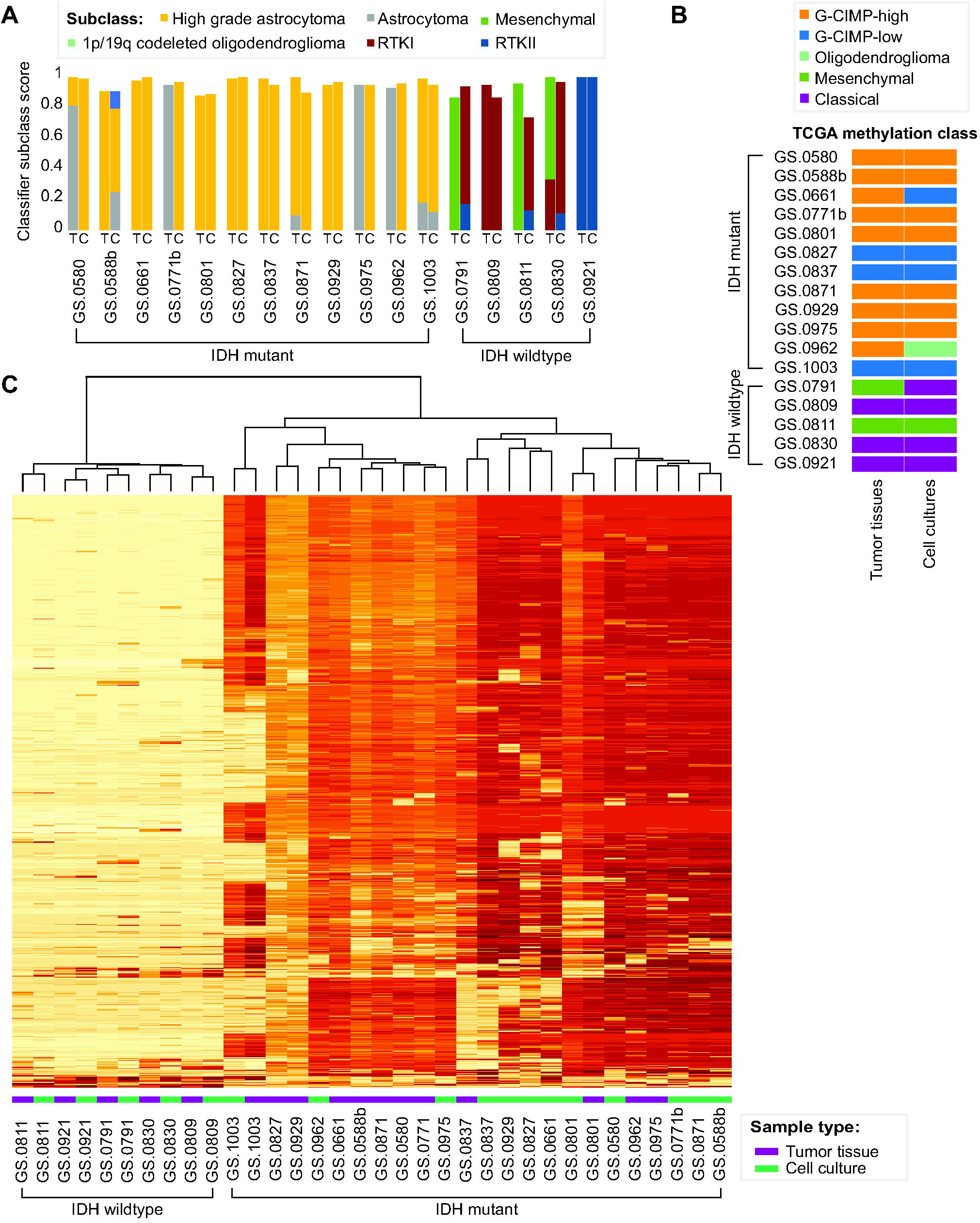
Global methylation profiles of IDHmt cell lines. **(A)** Bar graph of subclass scores from the global methylation-based CNS tumor classifier defined by Capper at al. The origin of the DNA sample, tumor tissue (T) or cell culture (C), is indicated. The Y-axis represents the methylation class family member classifier score (positive match score ≥ 0.5). **(B)** Representation of TCGA classification of tumor tissue and cell culture samples based on seven probes associated with G-CIMP high or G-CIMP low methylation class. **(C)** Heatmap of DNA methylation profile of five IDHwt tissue samples and cell cultures and twelve IDHmt tissue samples and cell cultures, based on the 1000 most variable methylated probes. The two main clusters are formed by IDHwt versus IDHmt cell cultures and tumor tissues.

TCGA-based classification of G-CIMP methylation profiles of our samples classified nine tumors as G-CIMP-high, while the remaining three gliomas were classified as G-CIMP-low (Fig. 2B).^23^ One of the G-CIMP-high tumors was G-CIMP-low in its corresponding cell culture, another switched to the oligodendroglioma subclass. Of the five IDHwt cultures all but one retained the G-CIMP classification. Unsupervised clustering of the 1000 most variably methylated CpG sites separated IDHwt tumors and cell cultures from IDHmt tumors and cell cultures (Fig. 2C), with IDHwt samples revealing lower overall methylation intensity. All IDHwt cultures cluster closest to their respective parental tumor sample. IDHmt tumor tissue samples tend to cluster together rather than with their matched cell culture; only four out of twelve IDHmt sample sets group together. Indeed, the primary (GS.0588b) and recurrent (GS.0871) tumor tissue samples derived from the same patient, cluster together, as do the derived cell cultures. This indicates longitudinal preservation of methylation profiles, both *in vitro* and *in situ*.

### Gene expression shows correlation between tumor and cell cultures

We performed RNA sequencing to evaluate the changes in expression profiles between tumor tissue and cell cultures. A PCA plot shows a very clear separation between tumor tissues and cell cultures in the first dimension (Fig. 3A). The second dimension separates IDHwt samples from IDHmt samples.

**Figure 3.**
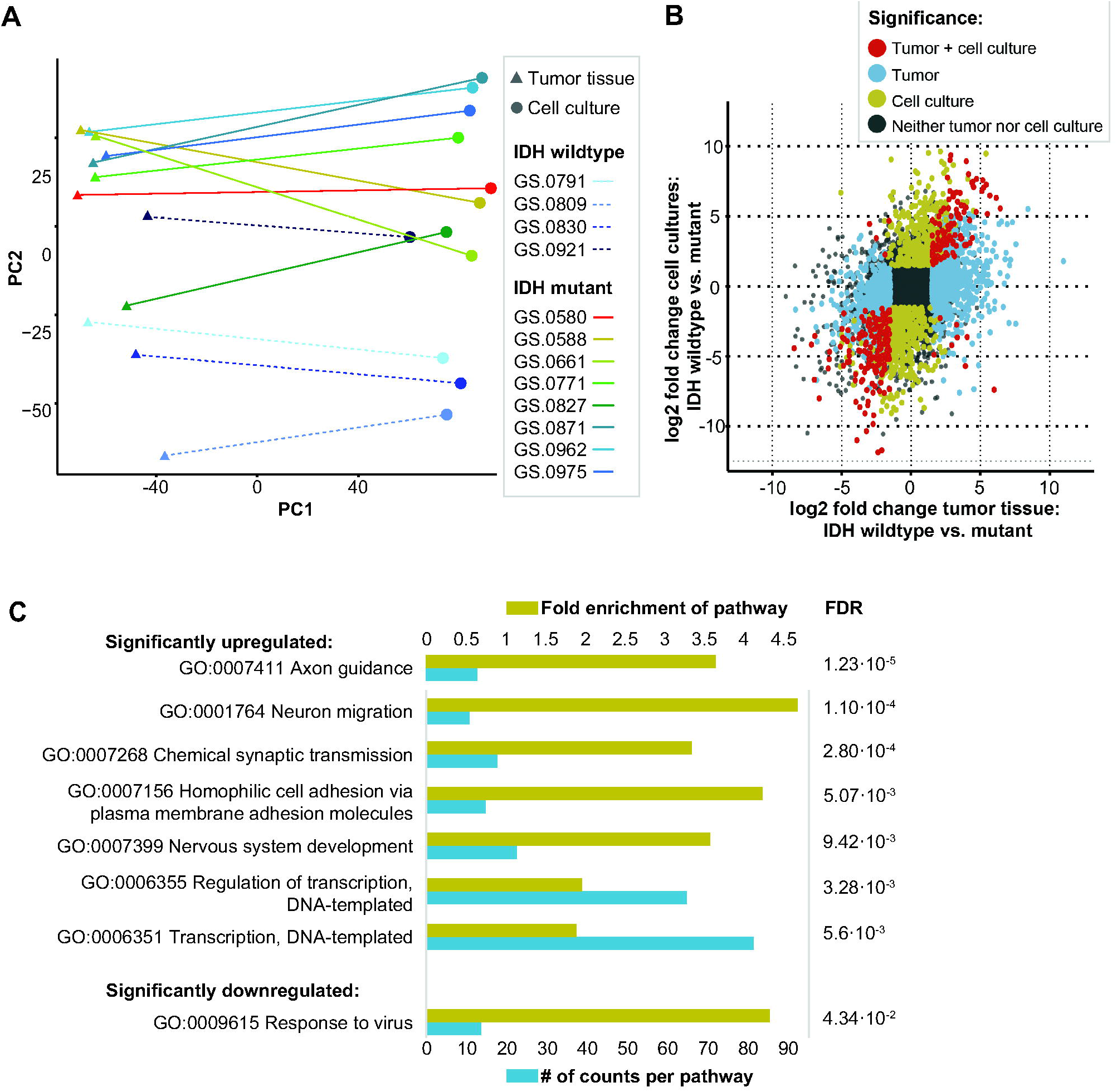
Correlation of transcriptomics of IDHmt cell cultures and parental tumor tissue. **(A)** Clustering of glioma cell cultures and tumor tissues based on RNA expression through principal component analysis. Components 1 and 2 are shown. **(B)** Scatterplot showing the correlation between the log2 fold change of IDHwt versus mutant in tumor tissues (X-axis) and IDHwt versus mutant in cell cultures (Y-axis). **(C)** Gene ontology pathway analysis of significantly upregulated or downregulated pathways in IDHmt versus IDHwt cell cultures (adjusted P-value < 0.05). The size of the blue bars represents the number of DEGs in that pathway (bottom x-axis), and yellow bars represents the fold enrichment (FE) of the pathway (top x-axis). The false discovery rate values are shown on the right side of the graph.

We correlated the log2 fold changes of IDHmt versus IDHwt tumor tissue with the log2 fold changes of IDHmt versus IDHwt cultures (adjusted p-value < 0.05, log2 [fold change] ≥ 0.58 for upregulated genes or ≤ −0.58 for downregulated genes) (Fig. 3B). There is an overall correlation, with a total of 161 genes significantly differentially expressed in both IDHmt tumor tissue and IDHmt cell cultures compared to their wild type counterparts. Gene ontology enrichment analysis identified upregulated differentially expressed genes involved in transcription, synaptic transmission and neural migration, and downregulated genes associated with response to virus infection (Fig 3C).

### The IDH mutation suppresses bioenergetic metabolism in cultured glioma

Oncometabolite D-2-HG has a myriad of effects on epigenetics and metabolism.^24^ Mitochondrial respiration was assessed by measuring extracellular oxygen consumption rate (OCR) and glycolysis through the acidification rate (ECAR), in six IDHmt and seven IDHwt cultures. Although basal OCR did not differ between IDHwt and IDHmt glioma cultures, mitochondrial reserve respiratory capacity was significantly reduced in IDH mutants (Fig. 4A). Moreover, mutant cells also displayed reduced ECAR, in both basal conditions and after inhibition of mitochondrial ATP synthase (adjusted P value < 0.05) (Fig. 4B). Canonical pathway analysis (KEGG) of RNA expression data, comparing IDHmt cell cultures with IDHwt cultures revealed that IDHmt cultures have downregulation of glycolysis and of other processes crucially involved in bioenergetics, such as pyruvate metabolism, the citric acid cycle, and nicotinate and nicotinamide metabolism, which all contribute to respiration substrates production (Table S4). Conversely, no differences were observed in the expression of genes of the electron transport chain complexes. Overall, evidence at transcription level is consistent with biochemical analyses showing reduced bioenergetic function in mutants and we conclude that IDHmt cell cultures have an intact electron transport chain but lower metabolite supply.

**Figure 4.**
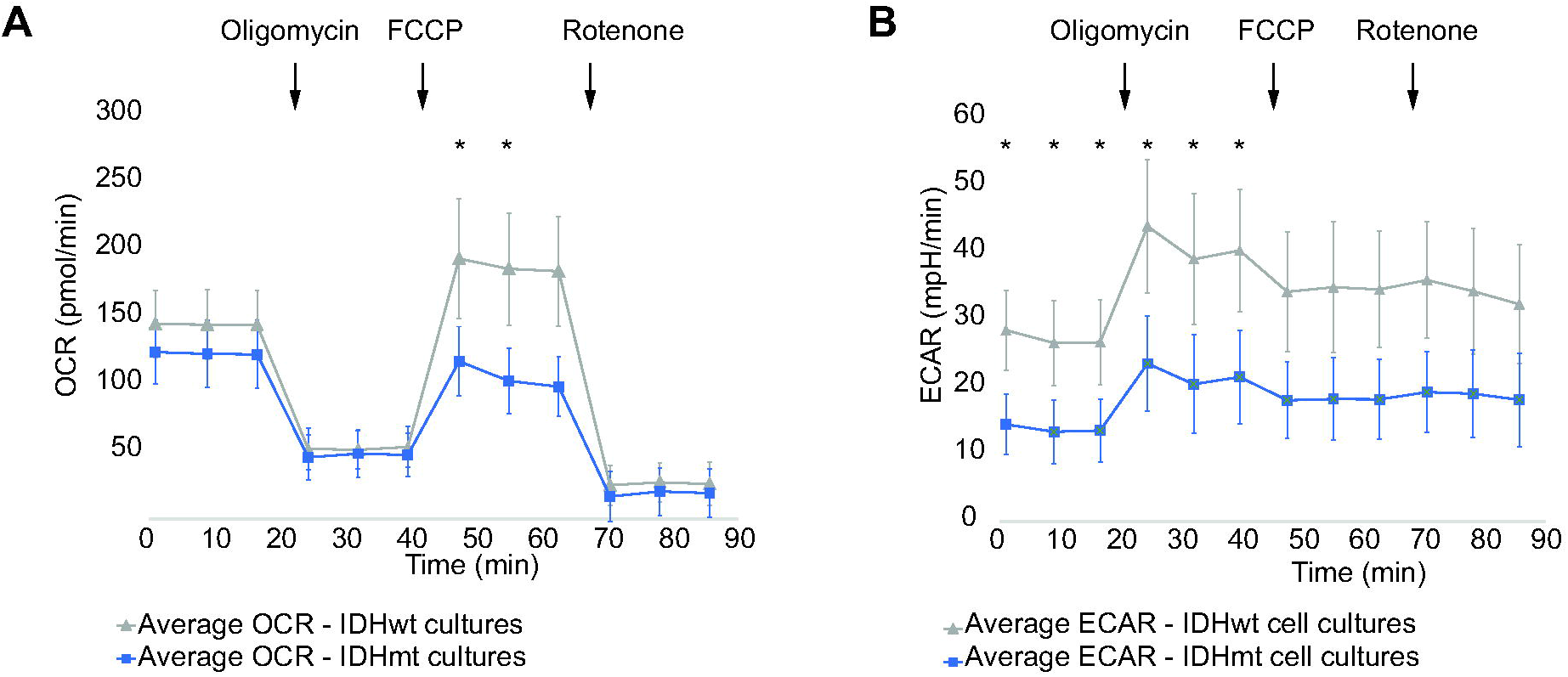
Live cell metabolic analysis of IDHmt glioma cultures. **(A)** Mean OCR values of IDHmt cultures (blue, N=6) and IDHwt cultures (grey, N=7). **(B)** Mean ECAR values of IDHmt cultures (blue, N=6) and IDHwt cultures (grey, N=7). Error bars represent the pooled standard deviations. Asterisks indicate significantly different mean OCR or ECAR values at the given timepoint (Independent Student’s T-test, P < 0.5). Error bars represent the SD. Ten technical replicates were tested for each culture.

### IDH mutant-specific inhibitors affect enzyme activity but not growth rate of IDHmt cells

Addition of IDH mutant-specific inhibitor AGI-5198 to the culture medium for seven days resulted in a dose-dependent decrease of D-2-HG levels in all IDHmt cell cultures (Fig. 5A). A concentration of 2.5 μM AGI-5198 was sufficient to reduce the D2HG levels with over 99%. The D2-HG levels of the two IDHwt cultures were below the detection limit. Despite this strong D-2-HG reduction, we did not observe any effect on cell viability. Similar results were obtained using an alternative inhibitor (Bay-1436032) (Fig. 5B). For both inhibitors, doses over 10 μM resulted in a decrease in cell viability, but this effect was also observed in IDHwt cultures, indicating more general toxicity. We also examined the effect on doubling times over three consecutive passages (Fig. 5C). Only IDHmt cell culture GS.0661 showed a significantly extended doubling time (p=0.02) in the presence of the 10 μM inhibitor, from 3.2 days for controls to 5.2 days in the presence of AGI-5198. These data suggest that the IDHmt cultures are not dependent on high D2-HG levels for viability or growth, at least not for the time spans investigated.

**Figure 5.**
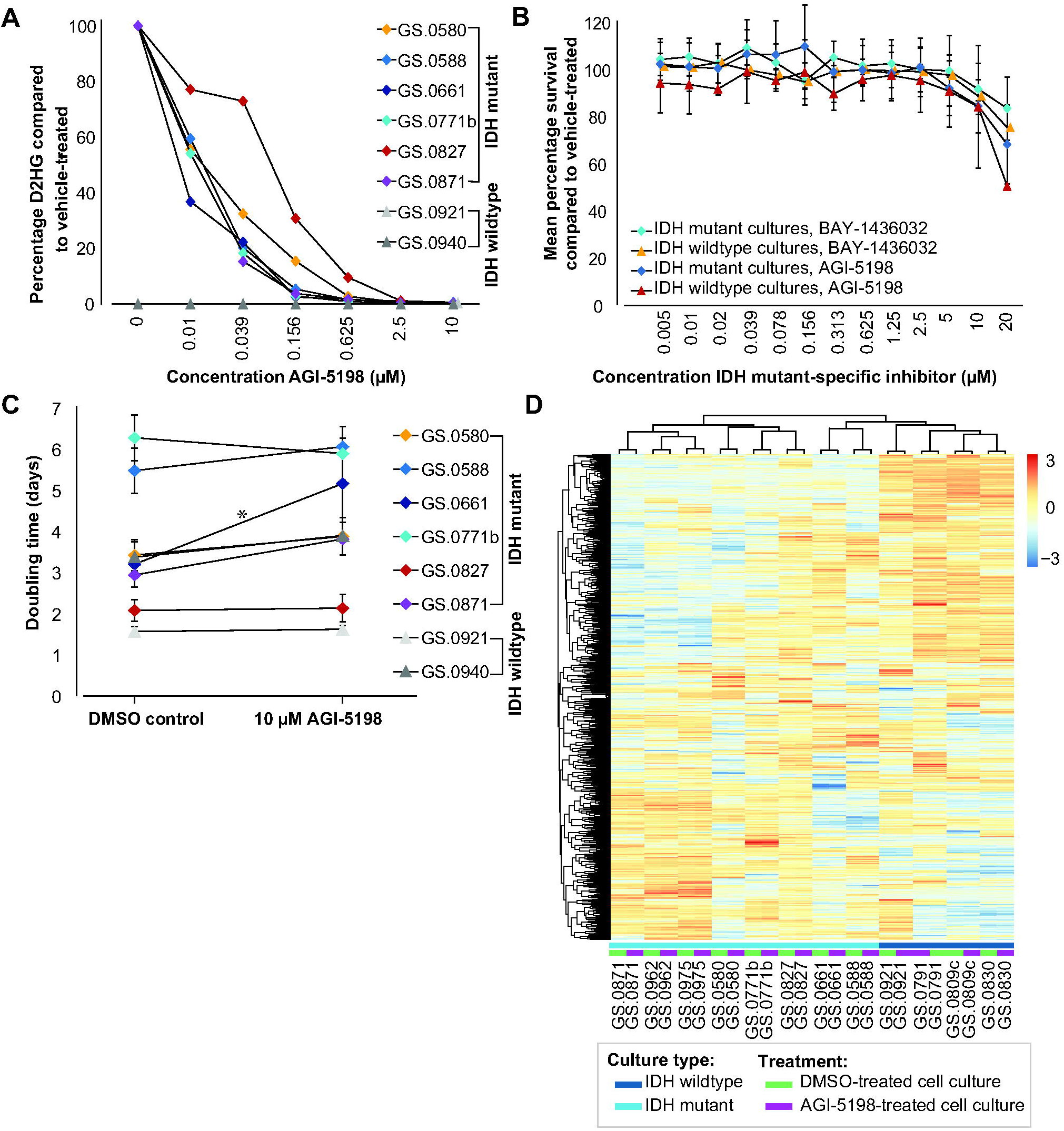
*in vitro* effect of IDH mutant-specific inhibitors on IDHmt glioma cultures. **(A)** Percentage of D2-HG in cell cultures treated with increasing concentrations of IDH-mutant specific inhibitor AGI-5198 compared to DMSO-controls. **(B)** Mean average survival of IDH-mutant and IDH-wildtype cultures treated with increasing concentrations of BAY-1438032 or AGI-5198. Data are presented as mean percentage survival ± SD. Each culture was tested with three technical replicates. **(C)** Mean doubling times of IDHmt (N=6) and IDHwt (N=2) cell cultures, cultured with and without 10 μM IDH mutant-specific inhibitor AGI-5198. Error bars represent standard deviations over three consecutive passages. (*P<0.05, paired T-test). Three technical replicates were used for each culture and time point. **(D)** Heatmap showing unsupervised cluster analysis of eight IDHmt glioma cultures and four IDHwt cultures, cultured with and without IDH mutant-specific inhibitor AGI-5198 for seven days. Clustering was performed on 1000 most variably expressed genes. The heatmap was scaled by row.

To address the effect of D2-HG on RNA expression, eight IDHmt and four IDHwt cultures were treated for seven days with the IDH mutant-specific inhibitor AGI-5198. Treatment with AGI-5198 does not result in a distinct expression signature and clustering analysis showed that treated and untreated cell culture counterparts remain nearest neighbors in all cases (Fig. 5D). In IDHmt glioma cultures, upon AGI-5198-treatment seven genes were significantly downregulated, and 17 upregulated (FDR < 0.00001, log2 [fold change] ≥ 1 or ≤ −1).

### Drug screening of 107 approved anti-cancer agents on IDHmt cell cultures

The seven first available IDHmt glioma cultures were screened for sensitivity to 107 FDA-approved oncology drugs. As an initial cut-off, we defined cultures as sensitive where IC_50_ values of the compounds were below reported C_max_ plasma values in patients.^25^ Nineteen compounds were identified that were effective in at least six of seven cultures (Fig. 6A). These compounds were grouped into nine drug subclasses and ranked within each class according to the CNS multi-parameter optimization (MPO) score, which predicts blood brain barrier (BBB) crossing based physicochemical properties (Table S5).^26^ However, compounds with low MPO score were still considered as possible candidates if enhanced or local delivery systems are in development for these drugs. Based on these data, we selected one compound from each subclass for further validation: gemcitabine, teniposide, daunorubicin, romidepsin, dactinomycin, regorafenib, and omacetaxine. We replaced the proteasome inhibitor bortezomib by marizomib, which has a similar mechanism of action and is currently under investigation in clinical GBM trials ^27,28^. For the taxanes we opted to include paclitaxel, as this compound is currently under investigation using focused ultrasound delivery (Table S5). ^29^

**Figure 6.**
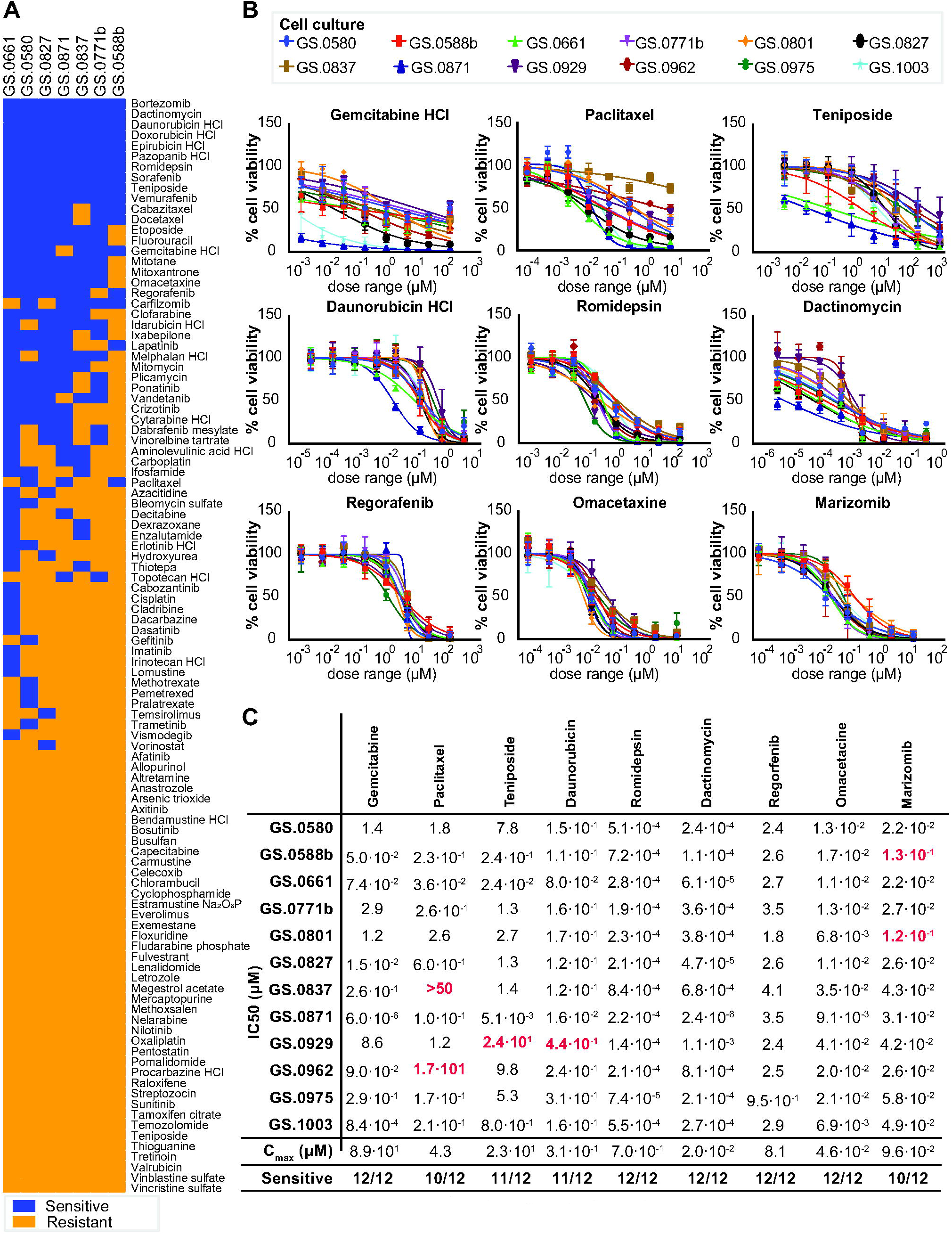
Drug screen with anti-cancer compounds in IDHmt glioma cultures. **(A)** Heatmap of response of 7 IDHmt glioma cultures to an FDA-approved anti-cancer drug set. Sensitive cell cultures (blue) were defined as having an IC50<C_max_ plasma, while resistant cultures had IC50 >C_max_. **(B)** Dose-response graphs of the anti-cancer compounds gemcitabine, paclitaxel, teniposide, daunorubicin, romidepsin, dactinomycin, regorafenib, omacetaxine and marizomib, on all twelve IDHmt cell cultures. The X-axes represent increasing concentrations of the respective compounds. Y-axes indicate the percentage cell viability compared to DMSO-treated controls. Error bars represent standard deviation of technical quadruplicates. At least two biological replicates were used for each cell culture. **(C)** IC50 values were calculated based on the dose-response curves shown in fig. 7C. All values represent the concentration in μM. Red numbers indicate IC50 > C_max_ and are labelled as resistant.

Using smaller-stepped dilution series, IC50 values of these drugs were determined on all twelve IDHmt cultures. This validation set revealed high overall sensitivity to each of the compounds (Fig. 6B), although a degree of intertumoral variability was observed. Notably, low-grade glioma culture GS.0962 showed most resistance to therapy. Mean IC50 values remained well-below reported plasma C_max_ values for most of the compounds, in particular for romidepsin and dactinomycin (Fig. 6C). These results present a set of available anti-cancer compounds that are effective against IDHmt glioma cells and which are either expected to cross the BBB, such as omacetaxin mepesuccinate and marizomib, or which are being developed for local or enhanced delivery, such as paclitaxel.

## Discussion

In this study, we present a panel of twelve cell cultures derived from 1p19q non-codeleted astrocytic tumors and an in-depth characterization of these cultures. Previously, we reported that EGF/FGF supplemented serum-free culture conditions do not support growth of IDHmt tumors.^6^ However, with adapted tumor dissociation and culture protocols, combined with careful monitoring of morphological features of the cells and in some cases isolation of astrocytic subpopulations, we demonstrate that such cultures can be established and maintained for a prolonged period of time. We identified morphological features that are an indicator of the stem-like nature of IDHmt glioma cells in adherent cell culture with brightfield microscopy, namely elongated cells with significantly smaller cell bodies, thin protrusions and bright edges. The relatively low success rate may be attributed to slow growth rate of most IDHmt gliomas, while culture success is based on expanding the fastest growing cells, which in some cases can result in stromal cells overcrowding the tumor cells in culture. Furthermore, culture conditions are very different from the local micro-environment within a tumor. This transition can be facilitated by the addition of growth factors to the culture medium, use of extracellular matrix coatings as we do in our protocol, or facilitating ex situ adaptation through serial xenografts.^14,30^

Homozygous loss of *CDKN2A/B* is a relatively common occurrence in gliomas and a predictor of poorer patient survival in IDHmt astrocytomas.^31^ Interestingly, without exception, we observed loss of *CDKN2A/B* in all IDHmt glioma cultures, whereas this was detected in only eight of the twelve the parental tumors. Loss of *CDKN2A/B* thus appears to be a prerequisite for successful culture establishment of IDHmt gliomas in our culture conditions. *CDKN2A/B* loss points towards a more aggressive phenotype and our cultures should therefore be considered as a representation of more advanced stages of astrocytomas, even when derived from low grade tumors.

Global methylation profiling is rapidly gaining momentum in the classification of central nervous system tumors.^17^ Using this classifier, all IDHmt astrocytoma cell cultures were assigned to high-grade astrocytomas (A_IDH_HG), which, on the genomic level, is in agreement with homozygous deletion of the *CDKN2A/B* locus. In a classifier as defined by the TCGA, such large differences were not identified and most cultures remained G-CIMP-high).

The functions of IDH mutations in metabolism are an important research focus, from a mechanistic point of view as well as to identify metabolic vulnerabilities that can be targeted for therapy.^21,32^ We measured mitochondrial respiration and glycolysis rate in our set of IDHmt cultures and correlated these parameters with gene expression data. This revealed that IDHmt glioma cultures have compromised bioenergetics and reduced mitochondrial reserve capacity, however, the electron transport chain remains unperturbed, suggesting that IDHmt cultures may have a lower metabolite supply. This is consistent with other reports that show reduced glycolysis in IDHmt patient and PDX tumors. Alternative carbon sources (such as glutamate, acetate and lactate) may supplement the bioenergetic fuel in IDHmt gliomas. ^10,33,34^ The availability of cell cultures with endogenous IDH mutation allows for further studies into these processes.

IDH mutations are early events in gliomagenesis which usually remain clonal as the tumor progresses.^35–37^ Occasional loss of mutant IDH in recurrent gliomas has been reported, suggesting that the presence of the mutation is not essential for tumor survival or tumor progression.^38^ Indeed, inhibition of the mutant enzyme and D2HG production did not decrease short-term viability, and only one cell culture showed a small decrease in growth rate when exposed for several passages. This contrasts with previous research that reports decreased growth rates *in vitro* and in xenografted mouse models.^39,40^ The source of this discrepancy may lie in the advanced grade of our cell cultures. If so, this may have important clinical implications as it suggests that patients with newly diagnosed low-grade glioma rather than secondary glioblastoma may be a better candidate for the use of IDH mutant-specific inhibitors. Alternatively, longer exposure to the inhibitor is required or effects of the inhibitor on the *in situ* tumor microenvironment play a role in its anti-glioma activity.

Drug screening of 107 FDA-approved anti-cancer compounds identified 19 compounds that showed effective killing at IC_50_ values well-below C_max_ plasma values. Although plasma C_max_ values do not accurately reflect drug concentrations in the tumor due to many factors including protein-binding, BBB penetration and the activity of drug efflux pumps, we used this value to discriminate between potentially effective compounds and those which are unlikely to reach effective concentrations in the brain. Interestingly, a number of the identified compounds have also been identified in previous *in vitro* compound screens on smaller sets of glial tumors.^41–45^ Several of the compounds identified in our screen have also been tested in clinical trials for glioma in the past (see table S5), but none of the compounds were tested specifically in IDH-mutant patients. The current developments in local or enhanced CNS delivery methods opens up new treatment possibilities for highly potent compounds that do not have favorable characteristics to pass the BBB, such as romidepsin and dactinomycin.^46,47^ Of particular interest are the compounds teniposide, omacetaxine, and marizomib, which are reported to cross the BBB and may offer candidates for systemic delivery ^48–50^.

In conclusion, we established a set of twelve patient-derived IDHmt glioma cell cultures from an astrocytoma background and different WHO grades. Based on the genetic, epigenetic, transcriptomic and metabolic similarity with the parental tumor, our IDHmt glioma cultures offer a versatile model system to test new therapeutic strategies and to advance research on this important subset of gliomas.

## Acknowledgements

We thank all patients for contributing to this research and the neurosurgeons involved for providing the tissue, Maurice de Wit, Alicia van der Ploeg, Judith van der Burg and Stanley Van for their contribution to data acquisition, and the NIH/NCI Developmental Therapeutics Program (http://dtp.cancer.gov) for providing the Oncology Drug Set.

