## Supplemental methods for "Generation, characterization and drug sensitivities of twelve patient-derived IDH1 mutant glioma cell cultures"

*Tumor processing and cell culture*

Resected tumor samples are collected in plain culture medium (DMEM, Gibco, Thermo Fisher Scientific) and kept at 4°C for a maximum of 4 hours until processing. Samples are processed according to a further optimized protocol based on Balvers et al^1^. Briefly, the tissue is first mechanically dissociated with a scalpel. All tumor fragments are then enzymatically dissociated with Collagenase A and DNAse (both from Roche). Red blood cells are removed by incubating the pellets with an erythrocyte lysis buffer. Single cells and tissue fragments are then separately resuspended in culture medium (DMEM-F12), supplemented with Penicillin/Streptomycin, B27, 20 ng/mL bFGF, 20 ng/mL EGF (all from Gibco, Thermo Fisher Scientific), 5 μg/mL Heparin (Alfa Aesar) and transferred to uncoated culture flasks. Tumor tissue fragments resected with a Cavitron Ultrasonic Surgical Aspirator (CUSA) device were also collected, of which the tissue fragments were scraped from the collection container and subsequently processed as normal tumor resection material. All cultures are kept at 37°C in a humidified incubator with 5% CO_2_.

After 5-8 days the cultures were transferred to a new flask coated with 1:100 diluted Cultrex PathClear Reduced Growth Factor BME (R&D Systems). Cultures were split when cultures reach >90% confluency or if there were multiple areas in the flasks of very high cell density. Cell cultures were considered successful if they could be passaged at least five times, in case of IDH mutant cell cultures the IDH mutation had to remain present. Colonies of cell types of interest were occasionally isolated by circling a colony with a P1000 pipette, gently scraping it off and aspirating it while monitoring this procedure through a brightfield microscope. The colony was then ejected into a new culture flask.

Cell cultures were tested for mycoplasma infection using the MycoAlert^TM^ PLUS Mycoplasma Detection Kit (Lonza). One culture tested positive at P1 (i.e., borderline detection) and was treated with Plasmocin, after which it was mycoplasma free.

*Culture growth assessment*

Growth rates and doubling times were assessed by seeding 2*10^5^ cells in a T75 flask pre-coated with1:100 diluted Cultrex PathClear Reduced Growth Factor BME (R&D Systems). When cultures were ready to be passaged (~90% confluency) they were split according to normal protocol and counted in triplicate using a hemocytometer. Then, 2·10^5^ cells were transferred to a new flask and the process was repeated at least two more times to evaluate differences over several passages.

To study the effect of inhibition of the mutant IDH enzyme on growth kinetics, we carried out the experiment as described above in the presence of 10 µM IDH-mutant-specific inhibitor AGI-5198 (Agios).

*Image acquisition*

Brightfield images were captured from live cells cultures on plastic T75 flasks coated with 1:100 diluted Cultrex PathClear Reduced Growth Factor BME (R&D Systems), with a Zeiss Axio Observer D1 Inverted Phase Contrast Fluorescence Microscope, HAL1 illuminator, and using a 10x Zeiss A-plan objective (421041-9910-000).

*Nucleic acid isolation*

Genomic DNA was extracted from pellets of cultured cells (passage numbers ranging from 0-12) and cryosections of snap frozen tumor material. Control, non-neoplastic DNA was isolated from leukocytes. All DNA was isolated with the DNeasy Blood & Tissue Kit (Qiagen) according to the manufacturer’s instructions. Extraction yields were determined with the Qubit dsDNA assay (Invitrogen, Thermo Fischer Scientific).

All RNA was extracted from pellets of cultured cells (passage numbers ranging from 8-14) and cryosections of snap frozen tumor material with the RNeasy Plus Mini Kit (Qiagen). Extraction yields were determined with the Qubit RNA HS assay (Invitrogen, Thermo Fischer Scientific).

*Analysis of IDH1 mutation status*

We subjected all cell culture candidates that could potentially harbor an *IDH* mutation to Sanger sequencing. We used genomic DNA for all sequencing experiments. For mutations in *IDH1* we used the primer set 5’-GTG GCA CGG TCT TCA GAG A-3’ and 5’- TTC ATA CCT TGC TTA ATG GGT GT- 3’.

*Exome sequencing*

Five hundred ng genomic DNA was fragmented to ~300 bp with the Kapa Hyperplus Library prep kit and unique dual indexed adapters libraries were generated. The libraries were amplified with six PCR cycles. The product size was checked on the Labchip GX (PerkinElmer) and concentrations were measured with Picogreen. Six samples were pooled equimolarly to a total amount of 1000 ng for exome capture. SepCap EZ Medexome probes (Roche) were added to each pool and incubated at 47°C for 72 hours. The exome of each pool was captured by the SeqCap wash kit and SeqCap pure capture bead kit (both from Roche). Another amplification PCR of 13 cycles were performed. The Illumina Novaseq platform was used to sequence 150 bases (paired end sequencing) to obtain 6GB per sample (coverage ~x).

Downstream analyses included demultiplexing (CASAVA software, Illumina) and subsequent alignment to human reference genome UCSC’s hg19 with the burrows-wheeler alignment tool ^2^. Alignments were sorted by Picard (http://broadinstitute.github.io/picard, v1.90) and then processed by GATK (Indel Realignment and Base-Quality Score Recalibration, v3.8;) ^3^. Lastly, PCR duplicates were marked by Picard, Mean Depth of Coverage was determined using GATK, and Freemix values were estimated through verifyBAMid ^4^. Samples that passed technical QC metrics were genotyped to gVCF level through GATKs HaplotypeCaller. Indels and SNVs were filtered separately using GATKs Variant-Quality Score Recalibration. Both the SNV and indel sets were annotated with ANNOVAR using freely available as well as customized databases ^5^. We analyzed the exome sequencing data to identify mutations annotated in the TCGA gene set ^6^. CopyNumbers were calculated using the CNVKit package ^7^.

*RNA sequencing*

RNA was isolated from cell cultures (+/- 5 µM AGI-5198) using the RNeasy kit (Qiagen). The Kapa mRNA Hyperprep Library prep kit (Roche) was used to prepare the RNA-seq library from 250ng total RNA per sample. From this, cDNA was generated to which unique dual indexed adapters were ligated. The cDNA library was amplified through PCR. Product size was checked on the Labchip GX (PerkinElmer). We performed paired-end sequencing of 2x100 with the Illumina Novaseq platform to obtain 8-10 GB per sample.

Reads were extracted from the raw sequencing data using CASAVA 1.8.2 (Illumina) and aligned to human reference genome UCSC’s hg19 with the STAR splice aware aligner (2.5.0c) using the gencode v19 transcriptome annotations as additional template ^8^. BAM files were then processed using various tools from the Picard Software Suite (v1.90; http://broadinstitute.github.io/picard), and tools from the Genome Analysis ToolKit (GATK, v3.8) ^3^. QC metrics were collected at various steps using Picard and evaluated, along with coverage metrics using GATK. Read counts per exon / gene were determined by the featureCounts function of the subread package (v1.4.6-p1) using the gencode v19 annotation as markers ^9^. Differential gene expression analysis and normalization was performed with DESeq2 and further visualizations of expression profiles used the DESeq2 VST transformed read counts ^10^. For differential expression analysis, only genes with on average 4 or more read counts across the tested samples were included.

*Principal component analysis and cluster analysis*

Principal component analysis was performed on the top 500 genes with the highest variance. We performed cluster analysis based on the 1000 most variably expressed genes using the pheatmap package in R, and scaled by row.

*Methylation Arrays*

Epigenome-wide DNA methylation levels were profiled by Infinium methylationEPIC beadchip arrays (Illumina) according to standard protocols. For classification according to methylation profile, data were uploaded to [www.molecularneuropathology.org](http://www.molecularneuropathology.org) and analyzed as reported elsewhere ^11^. The raw data were imported into R for statistical analysis using the Bioconductor *Minfi* package ^12^. TCGA subtypes were determined using the TCGAbiolinks Bioconductor package.

*Mitochondrial oxidative respiration assay*

We measured cellular oxidative respiration with the Seahorse XF24 Analyzer as described before ^13,14^. Briefly, glioma cell cultures were seeded on a Seahorse XF‐24 plate at a density of 4 · 10^4^ cells per well and grown overnight in our standard serum-free culture medium. We optimized the cell number based on an increased density curve of one wildtype and one mutant cell culture, from which we evaluated the optimal proportional response to Carbonyl cyanide-4-(trifluoromethoxy)phenylhydrazone (FCCP), a mitochondrial respiration uncoupler. The oxygen consumption rate (OCR) and extracellular acidification rate were recorded. One hour before the start of the experiment we changed the culture medium to unbuffered DMEM (XF Assay Medium; Agilent Technologies) supplemented with 10 mM glucose, 2 mM glutamin and 1 mM sodium pyruvate, with a pH adjusted to 7.4, and incubated at 37°C without CO_2_. After three baseline measurements, we measured cellular response after sequential injections of 1 μM oligomycin (Adenosine triphosphate (ATP) synthase inhibitor), 0.5 μM FCCP, and 1 μM antimycin (a complex III inhibitor). Each injection was followed by three measurements before the next injection was provided.

Basal respiration, mitochondrial ATP production, proton leakage, and maximal respiration rates were calculated with the Seahorse Wave software according to the manufacturer's instructions. For each cell culture, we used ten technical replicates.

*D/L-2-hydroxyglutarate measurements*

Intracellular D-2-HG and L-2-HG was measured in lysates of cell pellets after five days of culture. Samples were measured with LC-MS/MS as described by Struys *et al* ^15^. To study the effect of an IDH-mutant specific inhibitor, D-2-HG levels of cells treated for seven days with AGI-5198 were compared to D-2-HG levels of vehicle-treated cells.

*Drug screening*

For the drug screening studies, we tested 107 compounds from the Food and Drugs Administration Approved Oncology Drug Set II (National Cancer Institute) on the first seven IDH mutant glioma cultures that were available. The compounds Gemcitabine hydrochloride (Sigma-Aldrich), Paclitaxel (Sigma-Aldrich), Teniposide (Santa Cruz Biotechnology, Inc), Daunorubicin hydrochloride (SelleckChem), Romidepsin (MedChemExpress), Dactinomycin (BioViotica), Regorafenib (MedChemExpress), Omacetaxine mepesuccinate (Sigma-Aldrich), and Marizomib (Sigma-Aldrich), were selected for the validation studies. Initial screens were carried out in a 96-well format. Cells were seeded at a density of 1000 cells/well in wells pre-coated with 1:100 diluted Cultrex PathClear Reduced Growth Factor BME (R&D Systems). After 24 hours, serial dilutions of drugs were prepared and added to wells. For paclitaxel, dilutions were prepared in medium containing 0,1% bovine serum albumin (Sigma-Aldrich) to ensure solubility of this agent. After six days of exposure cell viability was measured with the ATP-based CellTiter GLO 2.0 kit (Promega) and luminescence was measured using the Infinite 200 reader (Tecan).

For the validation and expansion set experiments on the complete set of 12 IDH mutant cultures, screens were performed in 384-wells plates using an automated pipetting system (Hamilton) and cells were seeded in 4-plo at a density of 500 cells per well in wells pre-coated with Cultrex PathClear Reduced Growth Factor BME. Each of these 12 screens with 9 compounds was performed in 2-3 independent replicates. Serial dilutions of the selected compounds were added after 24 hours and viability was assessed by the CellTiter GLO 2.0 kit after five days.

*Effect of IDH-mutant specific inhibitors on viability*

We used IDH mutant specific inhibitors to assess the dependency of IDH mutant glioma cell cultures on their *IDH* mutations. Viability experiments were performed in 96-well formats with a seeding density of 1000 cells/well. We prepared two-fold dilution series of either AGI-5198 (Agios) or BAY-1436032 (Bayer), and added these to the designated wells. Viability was assessed after 5 and 8 days with the ATP-based CellTiter GLO 2.0 kit (Promega). Luminescence was measured using the Infinite 200 reader (Tecan).

*Pathway analysis*

We carried out an over-representation pathway analysis with the DAVID Bioinformatics Resources 6.8 (Huang DW, Sherman BT, Lempicki RA. Systematic and integrative analysis of large gene lists using DAVID Bioinformatics Resources. Nature Protoc. 2009;4(1):44-57). A pathway was identified as de-regulated with a false discovery rate (FDR) lower than 0.05. For each pathway was calculate a factor enrichment (FE) as the ratio of the DEGs and the number of genes annotates in this pathway. The FE is an indicator of the degree of enrichment. Additionally, pathway enrichment analysis was conducted via Gene Set Enrichment Analysis (GSEA) ^16^. Statistical significance of pathway enrichment score was ascertained by permutation testing with matched random gene sets. The proportion of false positives by multiple testing were controlled by FDR and a pathway was identified as de-regulated if the FDR was lower than 0.25 ^16^. Pathway information for GSEA was obtained from the Kyoto Encyclopedia of Genes and Genomes (KEGG) available at the Molecular Signatures Database (<http://www.broadinstitute.org/gsea/msigdb/index.jsp>).

*Statistical analysis*

The unpaired Student’s T-test was used for the comparison of two groups (statistical significance was defined as p < 0.05). To determine significance of contingency tables we used the Fisher exact test (statistical significance was defined as P < 0.05)

The IC50 values were calculated by applying a Nonlinear regression (curve-fit) and selecting the dose-response inhibition equation. Both IC50 and Area under the curve (AUC) calculations were done in GraphPad Prism version 9.
