## Supplementary material for "Generation, characterization and drug sensitivities of twelve patient-derived IDH1 mutant glioma cell cultures": Table S1

**Table S1. Clinical characteristics of patient samples from whom successful IDH mutant cell cultures were derived.**

| **GS#** | **PA** | **Grade** | **IDH mutation** | **Other notable mutations** | **Sex** | **Age at time of diagnosis (years)** | **Age at time of resection (years)** | **Overall survival (months)** | **Primary/ recurrent** | **PA primary tumor** | **Grade primary tumor** | **Tumor location** | **Therapy before surgery** | **Note** |
| --- | --- | --- | --- | --- | --- | --- | --- | --- | --- | --- | --- | --- | --- | --- |
| GS.0580 | Glioblastoma | IV | IDH1 R132H | TP53 | M | 31 | 40 | 122 | Recurrent | Diffuse astrocytoma | II | Right frontal | Stupp | NA |
| GS.0588 | Diffuse astrocytoma | II | IDH1 R132H | TP53, ATRX | F | 25 | 25 | 45 | Primary | NA | NA | Right frontal | NA | NA |
| GS.0661 | Glioblastoma | IV | IDH1 R132H | TP53 | F | 41 | 31 | 106 | Recurrent | Diffuse astrocytoma | II | Left frontal | Stupp | NA |
| GS.0771 | Glioblastoma | IV | IDH1 R132H | TP53, ATRX | F | 58 | 58 | NA | Primary | NA | NA | Right frontal | NA | NA |
| GS.0801 | Glioblastoma | IV | IDH1 R132H | NA | M | 46 | 51 | 53 | Recurrent | Diffuse astrocytoma | II | Left fronto-temporal | Stupp | NA |
| GS.0827 | Glioblastoma | IV | IDH1 R132H | TP53 | M | 30 | 31 | 20 | Recurrent | Diffuse astrocytoma | II | Right medio-frontal | Stupp | NA |
| GS.0837 | Glioblastoma | IV | IDH1 R132H | TP53 | F | 38 | 40 | 27 | Recurrent | Glioblastoma | IV | Left frontal | Stupp | NA |
| GS.0871 | Glioblastoma | IV | IDH1 R132H | TP53 | F | 25 | 28 | 45 | Recurrent (of GS588) | Diffuse astrocytoma | II | Right frontal | Radio-therapy, PVC, TMZ | Recurrent of GS588 |
| GS.0929 | Glioblastoma | IV | IDH1  R132H | TP53,  ATRX | M | 20 | 29 | 98 | Recurrent | Oligo-astrocytoma | II | Left frontal | Stupp, PVC | NA |
| GS.0962 | Diffuse astrocytoma | II | IDH1 R132H | TP53 | F | 47 | 47 | NA | Primary | NA | NA | Left frontal | NA | NA |
| GS.0975 | Anaplastic astrocytoma | III | IDH1 R132H | TP53, ATRX | M | 38 | 44 | NA | Recurrent | Diffuse astrocytoma | II | Right frontal | Stupp | NA |
| GS.1003 | Glioblastoma | IV | IDH1 R132H | TP53, ATRX | M | 33 | 46 | NA | Recurrent | Diffuse astrocytoma | II | Right temporal | NA | Date diagnosis based on first scan due to initial wait-and-see approach |
