## Supplementary material for "Generation, characterization and drug sensitivities of twelve patient-derived IDH1 mutant glioma cell cultures": Table S2

**Table S2. Characteristics and experimental use of IDH mutant cell cultures**

| **GS#** | **PA** | **IDHmt** | **WES** | **Growth speed** | **Global methy-lation array** | **RNA seq** | **Seahorse** | **D2HG/ L2HG ratio** | **Inhibitor BAY-146032 viability** | **Inhibitor AGI-5198 viability** | **Inhibitor AGI-5198 D2-HG** | **Inhibitor AGI-5198 doubling time** | **Initial drug screen** | **Validation drug screen** |
| --- | --- | --- | --- | --- | --- | --- | --- | --- | --- | --- | --- | --- | --- | --- |
| GS.0580 | Glioblastoma | IDH1 R132H | Yes | Yes | Yes | Yes | Yes | Yes | Yes | Yes | Yes | Yes | Yes | Yes |
| GS.0588 | Diffuse astrocytoma | IDH1 R132H | Yes | Yes | Yes | Yes | No | Yes | Yes | Yes | Yes | Yes | Yes | Yes |
| GS.0661 | Glioblastoma | IDH1 R132H | Yes | Yes | Yes | Yes | No | Yes | Yes | Yes | Yes | Yes | Yes | Yes |
| GS.0771 | Glioblastoma | IDH1 R132H | Yes | Yes | Yes | Yes | Yes | Yes | Yes | Yes | Yes | Yes | Yes | Yes |
| GS.0801 | Glioblastoma | IDH1 R132H | No | Yes | Yes | No | Yes | Yes | Yes | No | No | No | No | Yes |
| GS.0827 | Glioblastoma | IDH1 R132H | Yes | Yes | Yes | Yes | Yes | Yes | Yes | Yes | Yes | Yes | Yes | Yes |
| GS.0837 | Glioblastoma | IDH1 R132H | Yes | Yes | Yes | No | No | Yes | No | Yes | No | No | Yes | Yes |
| GS.0871 | Glioblastoma | IDH1 R132H | Yes | Yes | Yes | Yes | Yes | Yes | Yes | Yes | No | Yes | No | Yes |
| GS.0929 | Glioblastoma | IDH1  R132H | No | Yes | Yes | No | No | Yes | No | Yes | No | No | No | Yes |
| GS.0962 | Diffuse astrocytoma | IDH1 R132H | No | Yes | Yes | Yes | No | Yes | Yes | No | No | No | No | Yes |
| GS.0975 | Anaplastic astrocytoma | IDH1 R132H | No | Yes | Yes | Yes | Yes | Yes | Yes | Yes | No | No | No | Yes |
| GS.1003 | Glioblastoma | IDH1 R132H | No | Yes | Yes | No | Yes | Yes | Yes | Yes | No | No | No | Yes |
