## Supplementary material for "Generation, characterization and drug sensitivities of twelve patient-derived IDH1 mutant glioma cell cultures": Table S3

**Table S3 | Morphological characteristics of astrocytoma-like cells and fibroblast-like cells**

|  | Astrocytoma-like | Fibroblast-like |
| --- | --- | --- |
| Cell area (μm^2^) | 1646.78 ± 1111 | 6112 ± 4235 |
| Cell intensity | 1815.43 ± 1281 | 1263.20 ± 635 |
| Nucleus area (μm^2^) | 198.22 ± 86 | 300.27 ± 153 |
| Nucleus roundness | 0.86 ± 0.10 | 0.88 ± 0.12 |
| Nucleus intensity | 1926.03 ± 847 | 1018.76 ± 697 |
