## Supplementary material for "Generation, characterization and drug sensitivities of twelve patient-derived IDH1 mutant glioma cell cultures": Table S4

**Table S4. KEGG pathway analysis of RNA sequencing data of eight IDH mutant and four IDH wildtype glioma cell cultures.** De-regulated pathways were identified by GSEA when the transcriptome of IDH mutant cell cultures were compared with the transcriptome of IDH wild type cell cultures. This table relates to figure 5. **(A)** Up-regulated gene sets. **(B)** The top 20 down regulated gene sets. The oxidative phosphorylation pathway was not de-regulated, this pathway in enrichment with proteins of electron transport chain complexes. The deregulated pathways were ranked (RANK column) according to the FDR and the Normalized Enrichment Score (NES), which indicates the degree to which a gene set is overrepresented at the top or bottom of the pre-ranked list of genes. Size means the number of genes in the gene set.

**A.**

| Rank | Gene sets upregulated in idh-mutant vs idh-wildtype cell cultures | Size | NES | FDR q-val |
| --- | --- | --- | --- | --- |
| 1 | KEGG_TGF_BETA_SIGNALING_PATHWAY | 81 | 1.6 | 0.247 |
| 2 | KEGG_DNA_REPLICATION | 35 | 1.53 | 0.227 |
| 3 | KEGG_TASTE_TRANSDUCTION | 29 | 1.5 | 0.2 |

**B.**

| Rank | Gene sets downregulated in idh-mutant vs idh-wildtype cell cultures | Size | NES | FDR q-val |
| --- | --- | --- | --- | --- |
| 1 | KEGG_CELL_ADHESION_MOLECULES_CAMS | 99 | -1.72 | 0.16 |
| 2 | KEGG_REGULATION_OF_AUTOPHAGY | 18 | -1.71 | 0.094 |
| 3 | KEGG_TOLL_LIKE_RECEPTOR_SIGNALING_PATHWAY | 69 | -1.68 | 0.087 |
| 4 | KEGG_APOPTOSIS | 75 | -1.62 | 0.139 |
| 5 | KEGG_PATHOGENIC_ESCHERICHIA_COLI_INFECTION | 51 | -1.6 | 0.129 |
| 6 | **KEGG_NICOTINATE_AND_NICOTINAMIDE_METABOLISM** | **22** | **-1.59** | **0.126** |
| 7 | **KEGG_PYRUVATE_METABOLISM** | **34** | **-1.59** | **0.114** |
| 8 | KEGG_JAK_STAT_SIGNALING_PATHWAY | 98 | -1.55 | 0.144 |
| 9 | **KEGG_GLUTATHIONE_METABOLISM** | **40** | **-1.53** | **0.152** |
| 10 | KEGG_INSULIN_SIGNALING_PATHWAY | 122 | -1.52 | 0.159 |
| 11 | KEGG_HYPERTROPHIC_CARDIOMYOPATHY_HCM | 68 | -1.52 | 0.145 |
| 12 | KEGG_NATURAL_KILLER_CELL_MEDIATED_CYTOTOXICITY | 83 | -1.51 | 0.137 |
| 13 | KEGG_RIG_I_LIKE_RECEPTOR_SIGNALING_PATHWAY | 49 | -1.51 | 0.13 |
| 14 | **KEGG_GLYCINE_SERINE_AND_THREONINE_METABOLISM** | **27** | **-1.5** | **0.133** |
| 15 | KEGG_PROTEASOME | 40 | -1.5 | 0.129 |
| 16 | **KEGG_CITRATE_CYCLE_TCA_CYCLE** | **29** | **-1.48** | **0.148** |
| 17 | **KEGG_PENTOSE_PHOSPHATE_PATHWAY** | **21** | **-1.47** | **0.141** |
| 18 | **KEGG_GLYCOLYSIS_GLUCONEOGENESIS** | **48** | **-1.47** | **0.139** |
| 19 | **KEGG_ETHER_LIPID_METABOLISM** | **23** | **-1.46** | **0.139** |
| 20 | KEGG_HEMATOPOIETIC_CELL_LINEAGE | 54 | -1.45 | 0.145 |
| - | - | - | - | - |
| 56 | *KEGG_OXIDATIVE_PHOSPHORYLATION* | *108* | *-1.19* | *1* |
