## Supplementary material for "Generation, characterization and drug sensitivities of twelve patient-derived IDH1 mutant glioma cell cultures": Table S5

**Table S5. Anti-cancer compounds selected for validation on additional IDH-mutant cell cultures**. List of compounds to which at least six out of seven tested cultures were sensitive (IC50 < C_max_) in the drug screen. This table relates to figure 7. Paclitaxel (italic in the table, drug class taxanes) was not part of the best scoring compounds in the initial screen, but was chosen to represent this drug class in the validation set, as it is currently investigated in a phase I/II trial for glioblastoma with enhanced delivery methods. Marizomib (italic in the table), was not part of the initial screen but was selected to replace bortezomib as it is under clinical investigation for glioblastoma.

| **Drug class** | **Compounds** | **Mechanism of action (subclass)** | **CNS MPO-score**  **(Wager et al., 2010)** | **Protein binding**  **(DrugBank.com, 2020)** | **BBB-crossing** | **Enhanced delivery approaches** | **Previously tested in glioma patients** | **Selected for validation** |
| --- | --- | --- | --- | --- | --- | --- | --- | --- |
| Anti-metabolites | Fluorouracil | DNA synthesis inhibition | 5.5 | 8-12% | No (in vivo)  (Zhang et al., 2015) | Biodegradable microspheres (clinical)  (Menei et al 2005) |  | No |
|  | Gemcitabine hydrochloride | DNA synthesis inhibition | 3.95 | Negligible | No (clinical)  (Bernardi et al., 2008) | Convection enhanced delivery (clinical)  (Murad et al., 2007) | (Bastiancich et al., 2018) | Yes |
| Taxanes | Docetaxel | Microtubule inhibitor | 3.75 | 97% | Very limited  (ten Tije et al., 2004) | Polymeric micelles (in vivo)  (Zhang et al., 2013) |  | No |
|  | Cabazitaxel | Microtubule inhibitor | 2.75 | 82% | Yes (in vivo)  (Cisternino et al., 2003) | Ultrasound-based BBB-opening (in vivo)  (Sulheim et al., 2019) |  | No |
|  | *Paclitaxel* | Microtubule inhibitor | 1.5 | 89-98% | Very limited  (Fellner et al., 2002) | Ultrasound-based BBB-opening (clinical) (NCT04528680, 2020) | (Lidar et al., 2004) | Yes |
| Podophyllotoxins | Teniposide | Topoisomerase-II inhibition, induces single and double strand DNA breaks | 3.25 | NA | Yes (clinical)  (Philip A. Pizzo, 2011) | NA |  | Yes |
|  | Etoposide | Topoisomerase-II inhibition, induces single and double strand DNA breaks | 3.25 | 97% | Very limited  (Stewart et al., 1984) | Thermo-Responsive Biodegradable Paste (in vivo)  (Smith et al., 2019) | (Leonard and Wolff, 2013) | No |
| Anthracyclines | Daunorubicin hydrochloride | DNA intercalation and topoisomerase inhibition | 3.00 | 97% | Very limited (in vivo)  (Berg et al., 1999) | Liposome formulated (clinical)  (Albrecht et al., 2001) | (Fiorillo et al., 2004) | Yes |
|  | Doxorubicin hydrochloride | DNA intercalation and topoisomerase inhibition | 2.50 | 74-76% | No  (Sardi et al., 2013) | Monosialoganglioside  micelles (in vivo)  (Zou et al., 2017) |  | No |
|  | Epirubicin hydrochloride | DNA intercalation and topoisomerase inhibition | 2.50 | 77% | NA | Focused ultrasound/magnetic nanoparticle (in vivo)  (Liu et al., 2010) |  | No |
| HDAC inhibitor | Romidepsin | HDAC inhibition | 2.70 | 92-94% | Very limited (in vivo)  (Hiranaka et al., 2018) | NA | (Iwamoto et al., 2011) | Yes |
| Antineoplastic antibiotic | Mitoxantrone | DNA intercalation and topoisomerase-II inhibition | 2.78 | 78% | No  (Green et al., 1988) | NA |  | No |
|  | Dactinomycin | Transcription inhibitor | 2.00 |  | Very limited  AHFS Drug Information  Bethesda, MD. (2009) | NA |  | Yes |
| Tyrosine kinase inhibitors | Pazopanib hydrochloride | Multi-targeted tyrosine kinase inhibitor (VEGFR/PDGFR/KIT) | 3.67 | >99% | Yes (poorly)  (Iwamoto et al., 2010) | NA |  | No |
|  | Vemurafenib | BRAF/MEK/Erk pathway inhibitor | 2.06 | >99% | No  (Mittapalli et al., 2012) | NA |  | No |
|  | Sorafenib | Multi-targeted kinase inhibitor (VEGFR/PDGFR/KIT/BRAF) | 0.94 | >99% | No evidence | Lipid nanocapsules (in vivo)  (Clavreul et al., 2018) |  | No |
|  | Regorafenib | Multi-receptor kinase inhibitor (VEGFR/TIE) | 0.08 | >99% | Yes (clinical/CSF)  (Zeiner et al., 2019) | NA | (Lombardi et al., 2019) | Yes |
| Cephalotaxus alkaloids | Omacetaxine mepesuccinate | Protein synthesis inhibitor | 3.95 | 50% | Yes  (Savaraj et al., 1987) | NA | (Feun et al., 1990) | Yes |
| Proteasome inhibitors | *Marizomib* | Inhibition of 20S proteasome | 5.5 | NA | Yes  (Di et al., 2016; Millward et al., 2012) | NA |  | Yes |
|  | Bortezomib | Inhibition of 26S proteasome | 2.58 | 83% | Very limited  (Huehnchen et al., 2020; Yu et al., 2006) | NA |  | Replaced by Marizomib |
| Miscellaneous | Mitotane | Unclear | 2.9 | 6% | Very limited  (American Society of Health System Pharmacists, 2009) | NA |  | No (cytotoxicity) (Pape et al., 2018) |

**References Table S5**

Albrecht, K.W., de Witt Hamer, P.C., Leenstra, S., Bakker, P.J.M., Beijnen, J.H., Troost, D., Kaaijk, P., and Bosch, A.D. (2001). High Concentration of Daunorubicin and Daunorubicinol in Human Malignant Astrocytomas after Systemic Administration of Liposomal Daunorubicin. Journal of Neuro-Oncology *53*, 267-271.

Bastiancich, C., Bastiat, G., and Lagarce, F. (2018). Gemcitabine and glioblastoma: challenges and current perspectives. Drug Discov Today *23*, 416-423.

Berg, S.L., Reid, J., Godwin, K., Murry, D.J., Poplack, D.G., Balis, F.M., and Ames, M.M. (1999). Pharmacokinetics and cerebrospinal fluid penetration of daunorubicin, idarubicin, and their metabolites in the nonhuman primate model. J Pediatr Hematol Oncol *21*, 26-30.

Bernardi, R.J., Bomgaars, L., Fox, E., Balis, F.M., Egorin, M.J., Lagattuta, T.F., Aikin, A., Whitcomb, P., Renbarger, J., Lieberman, F.S.*, et al.* (2008). Phase I clinical trial of intrathecal gemcitabine in patients with neoplastic meningitis. Cancer Chemother Pharmacol *62*, 355-361.

Clavreul, A., Roger, E., Pourbaghi-Masouleh, M., Lemaire, L., Tétaud, C., and Menei, P. (2018). Development and characterization of sorafenib-loaded lipid nanocapsules for the treatment of glioblastoma. Drug Deliv *25*, 1756-1765.

Di, K., Lloyd, G.K., Abraham, V., Maclaren, A., Burrows, F.J., Desjardins, A., Trikha, M., and Bota, D.A. (2016). Marizomib activity as a single agent in malignant gliomas: ability to cross the blood-brain barrier. Neuro-Oncology *18*, 840-848.

DrugBank.com (2020). Protein binding.

Feun, L.G., Savaraj, N., Landy, H., Levin, H., and Lampidis, T. (1990). Phase II study of homoharringtonine in patients with recurrent primary malignant central nervous system tumors.  *9*, 159-163.

Fiorillo, A., Maggi, G., Greco, N., Migliorati, R., D'Amico, A., De Caro, M.D., Sabbatino, M.S., and Buffardi, F. (2004). Second-line chemotherapy with the association of liposomal daunorubicin, carboplatin and etoposide in children with recurrent malignant brain tumors. J Neurooncol *66*, 179-185.

Green, R.M., Stewart, D.J., Hugenholtz, H., Richard, M.T., Thibault, M., and Montpetit, V. (1988). Human central nervous system and plasma pharmacology of mitoxantrone. Journal of Neuro-Oncology *6*, 75-83.

Hiranaka, S., Tega, Y., Higuchi, K., Kurosawa, T., Deguchi, Y., Arata, M., Ito, A., Yoshida, M., Nagaoka, Y., and Sumiyoshi, T. (2018). Design, Synthesis, and Blood–Brain Barrier Transport Study of Pyrilamine Derivatives as Histone Deacetylase Inhibitors. ACS Medicinal Chemistry Letters *9*, 884-888.

Huehnchen, P., Springer, A., Kern, J., Kopp, U., Kohler, S., Alexander, T., Hiepe, F., Meisel, A., Boehmerle, W., and Endres, M. (2020). Bortezomib at therapeutic doses poorly passes the blood-brain-barrier and does not impair cognition. Brain Communications.

Iwamoto, F.M., Lamborn, K.R., Kuhn, J.G., Wen, P.Y., Alfred Yung, W.K., Gilbert, M.R., Chang, S.M., Lieberman, F.S., Prados, M.D., and Fine, H.A. (2011). A phase I/II trial of the histone deacetylase inhibitor romidepsin for adults with recurrent malignant glioma: North American Brain Tumor Consortium Study 03-03. Neuro-Oncology *13*, 509-516.

Iwamoto, F.M., Lamborn, K.R., Robins, H.I., Mehta, M.P., Chang, S.M., Butowski, N.A., Deangelis, L.M., Abrey, L.E., Zhang, W.T., Prados, M.D.*, et al.* (2010). Phase II trial of pazopanib (GW786034), an oral multi-targeted angiogenesis inhibitor, for adults with recurrent glioblastoma (North American Brain Tumor Consortium Study 06-02).  *12*, 855-861.

Leonard, A., and Wolff, J.E. (2013). Etoposide improves survival in high-grade glioma: a meta-analysis. Anticancer Res *33*, 3307-3315.

Lidar, Z., Mardor, Y., Jonas, T., Pfeffer, R., Faibel, M., Nass, D., Hadani, M., and Ram, Z. (2004). Convection-enhanced delivery of paclitaxel for the treatment of recurrent malignant glioma: a Phase I/II clinical study.  *100*, 472-479.

Liu, H.L., Hua, M.Y., Yang, H.W., Huang, C.Y., Chu, P.C., Wu, J.S., Tseng, I.C., Wang, J.J., Yen, T.C., Chen, P.Y.*, et al.* (2010). Magnetic resonance monitoring of focused ultrasound/magnetic nanoparticle targeting delivery of therapeutic agents to the brain. Proceedings of the National Academy of Sciences *107*, 15205-15210.

Lombardi, G., De Salvo, G.L., Brandes, A.A., Eoli, M., Rudà, R., Faedi, M., Lolli, I., Pace, A., Daniele, B., Pasqualetti, F.*, et al.* (2019). Regorafenib compared with lomustine in patients with relapsed glioblastoma (REGOMA): a multicentre, open-label, randomised, controlled, phase 2 trial. Lancet Oncol *20*, 110-119.

Millward, M., Price, T., Townsend, A., Sweeney, C., Spencer, A., Sukumaran, S., Longenecker, A., Lee, L., Lay, A., Sharma, G.*, et al.* (2012). Phase 1 clinical trial of the novel proteasome inhibitor marizomib with the histone deacetylase inhibitor vorinostat in patients with melanoma, pancreatic and lung cancer based on in vitro assessments of the combination. Invest New Drugs *30*, 2303-2317.

Murad, G.J., Walbridge, S., Morrison, P.F., Szerlip, N., Butman, J.A., Oldfield, E.H., and Lonser, R.R. (2007). Image-guided convection-enhanced delivery of gemcitabine to the brainstem. J Neurosurg *106*, 351-356.

NCT04528680 (2020). Ultrasound-based Blood-brain Barrier Opening and Albumin-bound Paclitaxel for Recurrent Glioblastoma (SC9/ABX) (Bethesda ).

Pape, E., Feliu, C., Yéléhé-Okouma, M., Colling, N., Djerada, Z., Gambier, N., Weryha, G., and Scala-Bertola, J. (2018). High-Dose Mitotane-Induced Encephalopathy in the Treatment of Adrenocortical Carcinoma. Oncologist *23*, 389-390.

Philip A. Pizzo, D.G.P. (2011). Principles and Practice of Pediatric Oncology, 6th edn (Philadelphia: Lippincott Williams & Wilkins).

Sardi, I., la Marca, G., Cardellicchio, S., Giunti, L., Malvagia, S., Genitori, L., Massimino, M., de Martino, M., and Giovannini, M.G. (2013). Pharmacological modulation of blood-brain barrier increases permeability of doxorubicin into the rat brain. Am J Cancer Res *3*, 424-432.

Savaraj, N., Feun, L.G., Lu, K., Leavens, M., Moser, R., Fields, W.S., and Loo, T.L. (1987). Central nervous system (CNS) penetration of homoharringtonine (HHT).  *5*, 77-81.

Smith, S.J., Tyler, B.M., Gould, T., Veal, G.J., Gorelick, N., Rowlinson, J., Serra, R., Ritchie, A., Berry, P., Otto, A.*, et al.* (2019). Overall Survival in Malignant Glioma Is Significantly Prolonged by Neurosurgical Delivery of Etoposide and Temozolomide from a Thermo-Responsive Biodegradable Paste. Clinical Cancer Research *25*, 5094-5106.

Stewart, D., Richard, M., Hugenholtz, H., Dennery, J., Belanger, R., Gerin-Lajoie, J., Montpetit, V., Nundy, D., Prior, J., and Hopkins, H. (1984). Penetration of VP-16 (etoposide) into human intracerebral and extracerebral tumors.  *2*.

Sulheim, E., Mørch, Y., Snipstad, S., Borgos, S.E., Miletic, H., Bjerkvig, R., Davies, C.D.L., and Åslund, A.K.O. (2019). Therapeutic Effect of Cabazitaxel and Blood-Brain Barrier opening in a Patient-Derived Glioblastoma Model. Nanotheranostics *3*, 103-112.

ten Tije, A.J., Loos, W.J., Zhao, M., Baker, S.D., Enting, R.H., van der Meulen, H., Verweij, J., and Sparreboom, A. (2004). Limited cerebrospinal fluid penetration of docetaxel. Anticancer Drugs *15*, 715-718.

Wager, T.T., Hou, X., Verhoest, P.R., and Villalobos, A. (2010). Moving beyond Rules: The Development of a Central Nervous System Multiparameter Optimization (CNS MPO) Approach To Enable Alignment of Druglike Properties. ACS Chemical Neuroscience *1*, 435-449.

Yu, L.J., Riordan, B., Hatsis, P., Brockman, A., Daniels, S., Stagliano, N., Finklestein, S., Ren, J., Milton, M., and Miwa, G. (2006). Study of brain and whole blood PK/PD of bortezomib in rat models. Journal of Clinical Oncology *24*, 12036-12036.

Zeiner, Kinzig, Divé, Maurer, Filipski, Harter, Senft, Bähr, Hattingen, Steinbach*, et al.* (2019). Regorafenib CSF Penetration, Efficacy, and MRI Patterns in Recurrent Malignant Glioma Patients. Journal of Clinical Medicine *8*, 2031.

Zhang, J., Zhang, L., Yan, Y., Li, S., Xie, L., Zhong, W., Lv, J., Zhang, X., Bai, Y., and Cheng, Z. (2015). Are Capecitabine and the Active Metabolite 5-FU CNS Penetrable to Treat Breast Cancer Brain Metastasis? Drug Metabolism and Disposition *43*, 411-417.

Zhang, Z., Wei, X., Zhang, X., and Lu, W. (2013). p-Hydroxybenzoic acid (p-HA) modified polymeric micelles for brain-targeted docetaxel delivery. Chinese Science Bulletin *58*, 2651-2656.

Zou, D., Wang, W., Lei, D., Yin, Y., Ren, P., Chen, J., Yin, T., Wang, B., Wang, G., and Wang, Y. (2017). Penetration of blood-brain barrier and antitumor activity and nerve repair in glioma by doxorubicin-loaded monosialoganglioside micelles system. International Journal of Nanomedicine *Volume 12*, 4879-4889.
