## Supplementary material for "Generation, characterization and drug sensitivities of twelve patient-derived IDH1 mutant glioma cell cultures": Legends supplementary figures

**Fig. S1. IDH mutant glioma cultures grow slower compared with IDH wildtype cultures, and faithfully produce D-2-HG in culture. (A)** Doubling time of IDH mutant glioma cultures (N=11) and IDH wild type cultures (N=12) over three consecutive passages. Error bars represent the standard deviation for these three timepoints. **(B)** Bar graph representing the D2-HG over L2-HG ratio of IDH mutant glioma cultures (N=8) and IDH wild type cultures (N=4), determined by LC-MS/MS from snap frozen cell pellets. Pellets were collected seven days after seeding. Cell cultures were between passage 8 and passage 14.

**Fig. S2. Copy number plots of glioma cell cultures show resemblance to those derived from their parental tumor tissue samples.** Copy number plots of IDH mutant (A-L) and IDH wild type (M-Q) tumor tissues and daughter cell cultures, retrieved from [www.molecularneuropathology.org](http://www.molecularneuropathology.org) and based on uploaded global methylation Infinium methylationEPIC beadchip array data.

**Fig. S3. Copy number plots of IDH mutant glioma GS.0771 and its daughter cell cultures GS.0771a (loss of mutant allele) and GS.0771b (mutant allele stably present in culture).**

**Fig. S4. IDH mutant glioma cultures have astrocytoma-like morphologies when visualised with brightfield microscopy of live cells. (A-L)** Brightfield images at 10x magnification of adherent IDH mutant cell cultures, showing inter-tumor heterogeneity in morphology but all typical astrocytoma-like characteristics: bright edges, small cell bodies and thin protrusions. **(M-O)** Brightfield images at 10x magnification of adherent cell cultures derived from IDH mutant gliomas, but upon evaluation with loss of the mutant allele and fibroblast-like phenotype.
