## Supplementary figures and images for "Generation, characterization and drug sensitivities of twelve patient-derived IDH1 mutant glioma cell cultures"

### Fig. S1

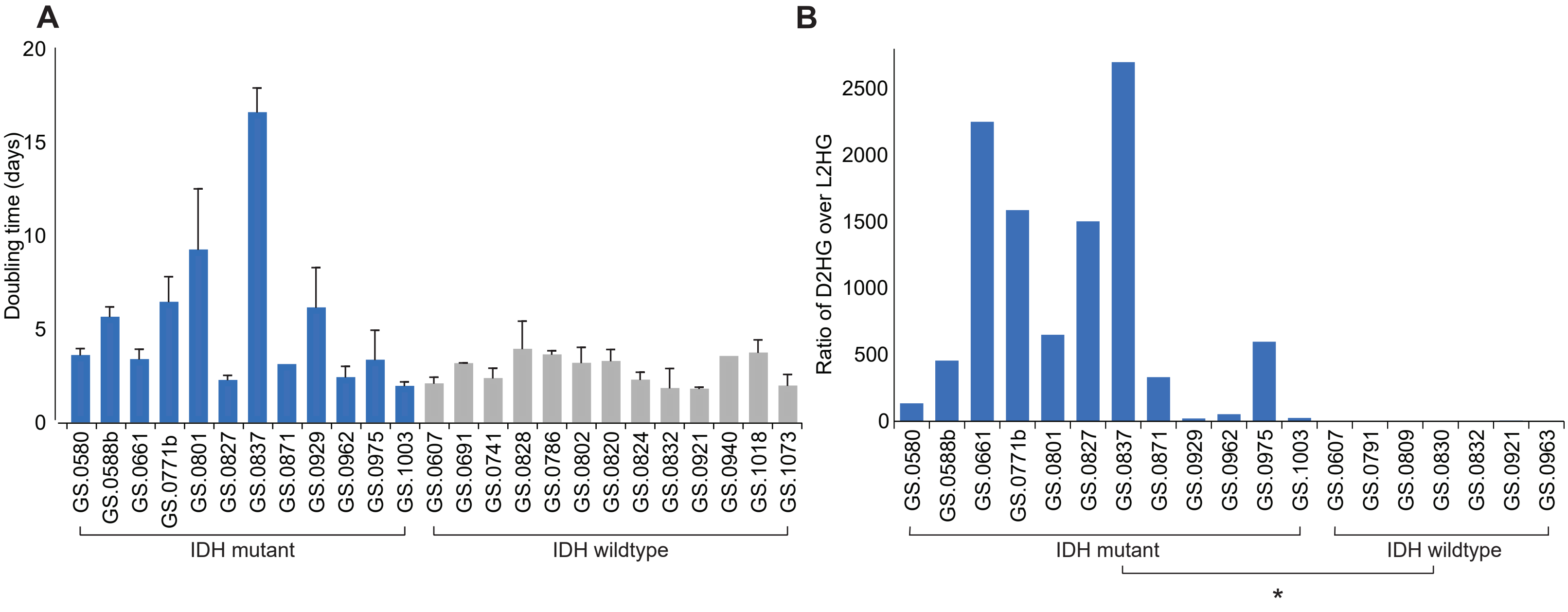


**Fig. S1 | Doubling time and D2-HG production of glioma cell cultures**

### Fig. S3

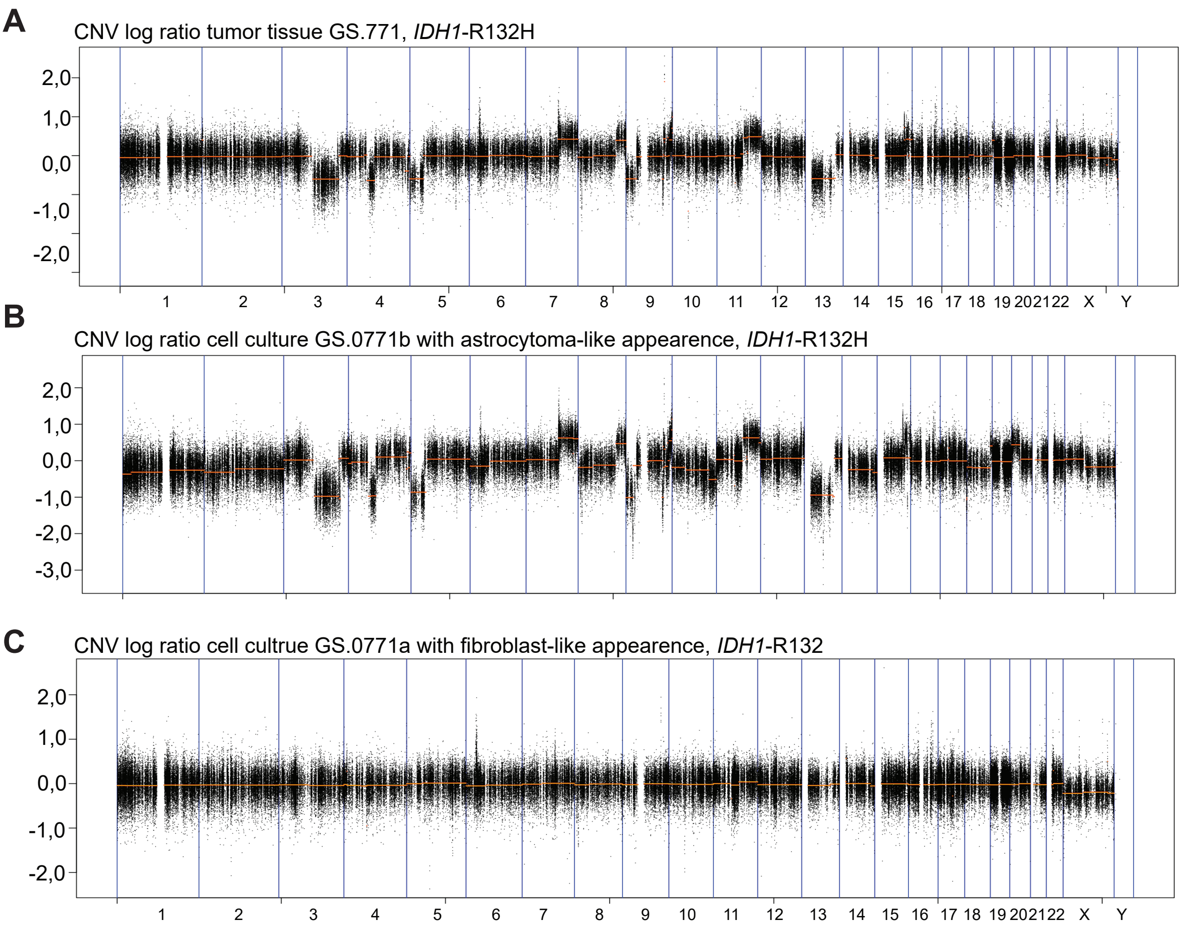


**Fig. S3 | Copy number profiles of IDHmt glioma GS.0771 and two daughter cell cultures**
