## Supplementary material for "Generation, characterization and drug sensitivities of twelve patient-derived IDH1 mutant glioma cell cultures": Fig. S2

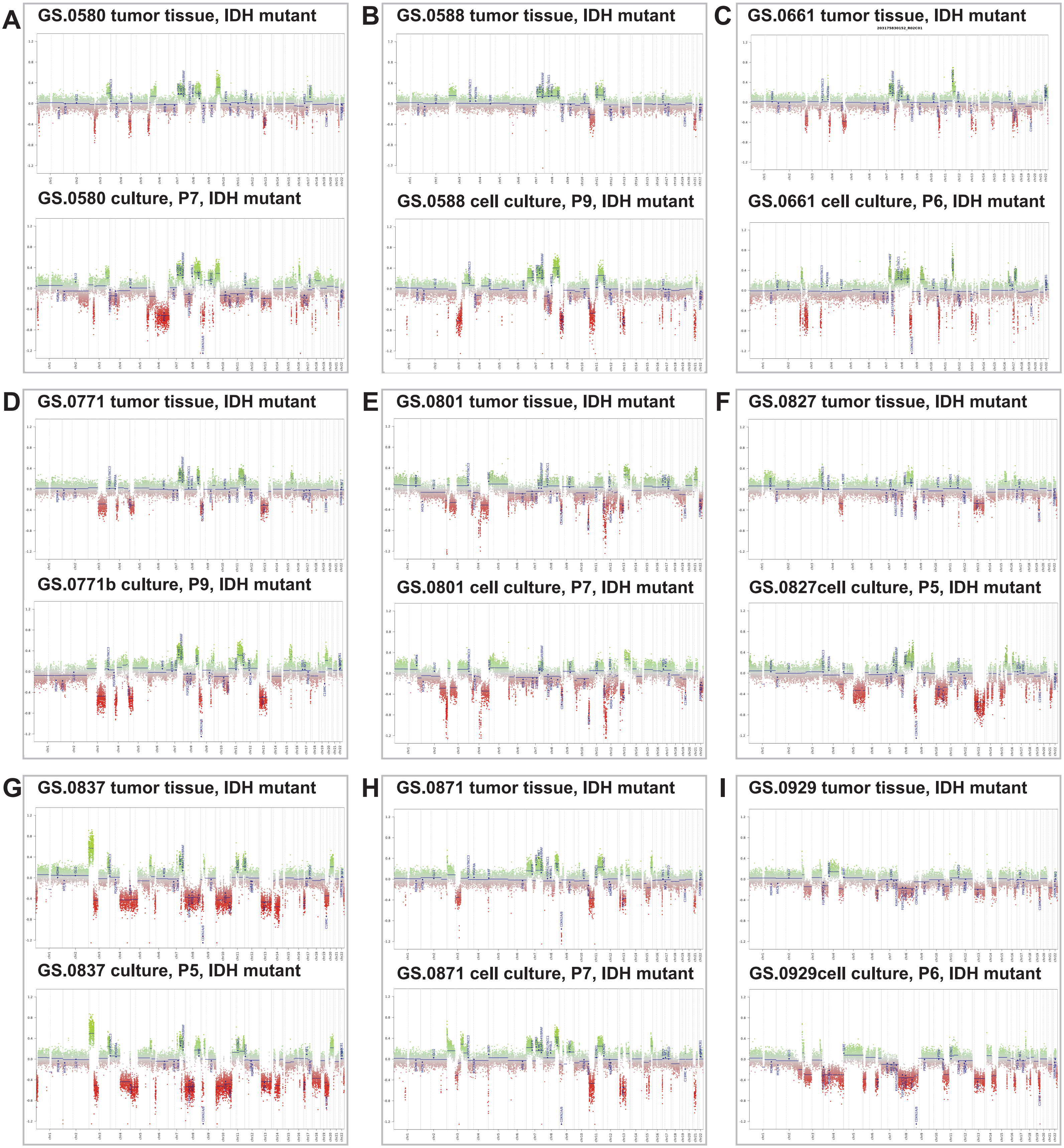


**Fig. S2 | Copy number plot of matched tissue/tumor samples**

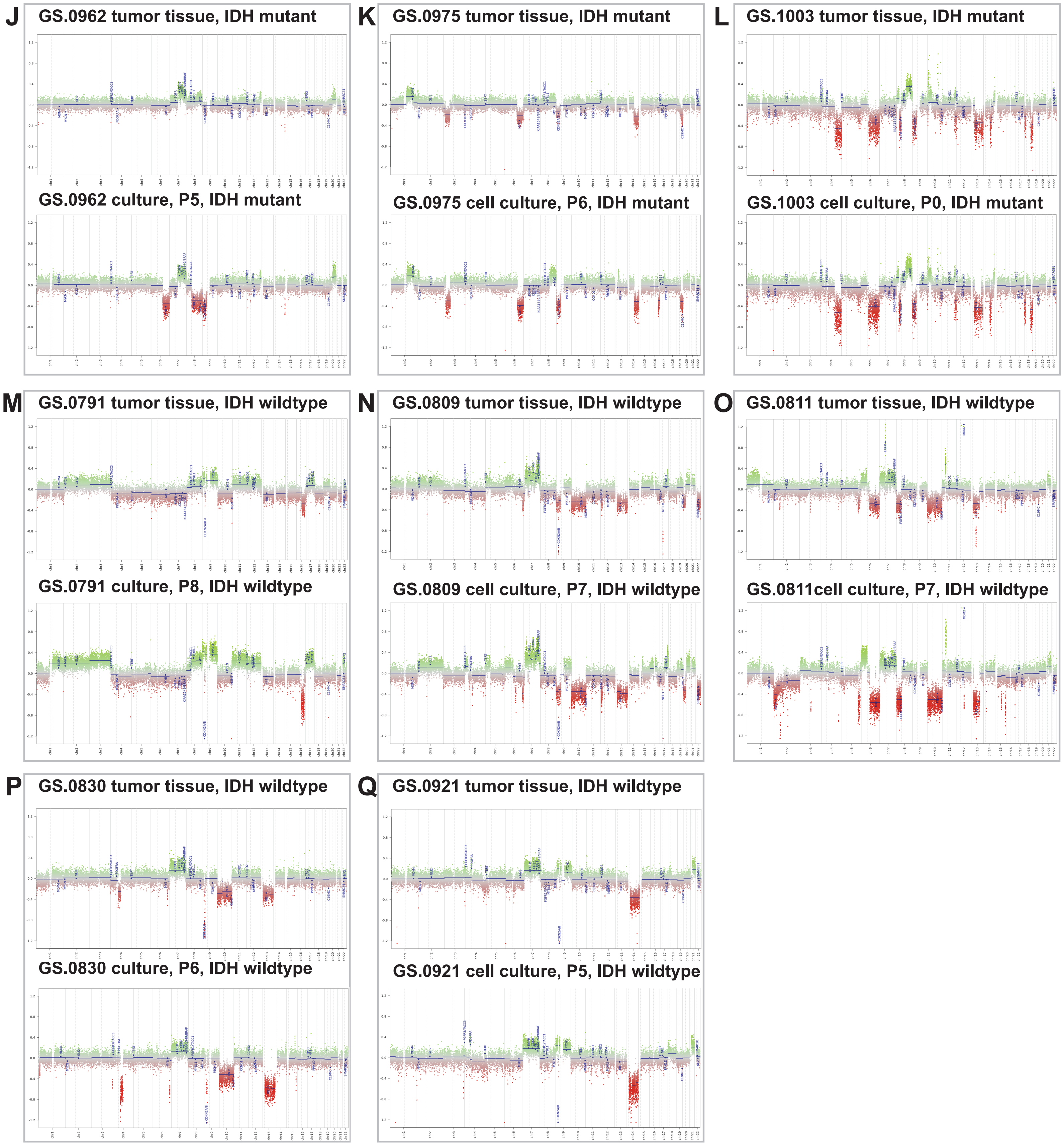


**Fig. S2 (continued) | Copy number plot of matched tissue/tumor samples**
