## Supplementary material for "Generation, characterization and drug sensitivities of twelve patient-derived IDH1 mutant glioma cell cultures": Fig. S4

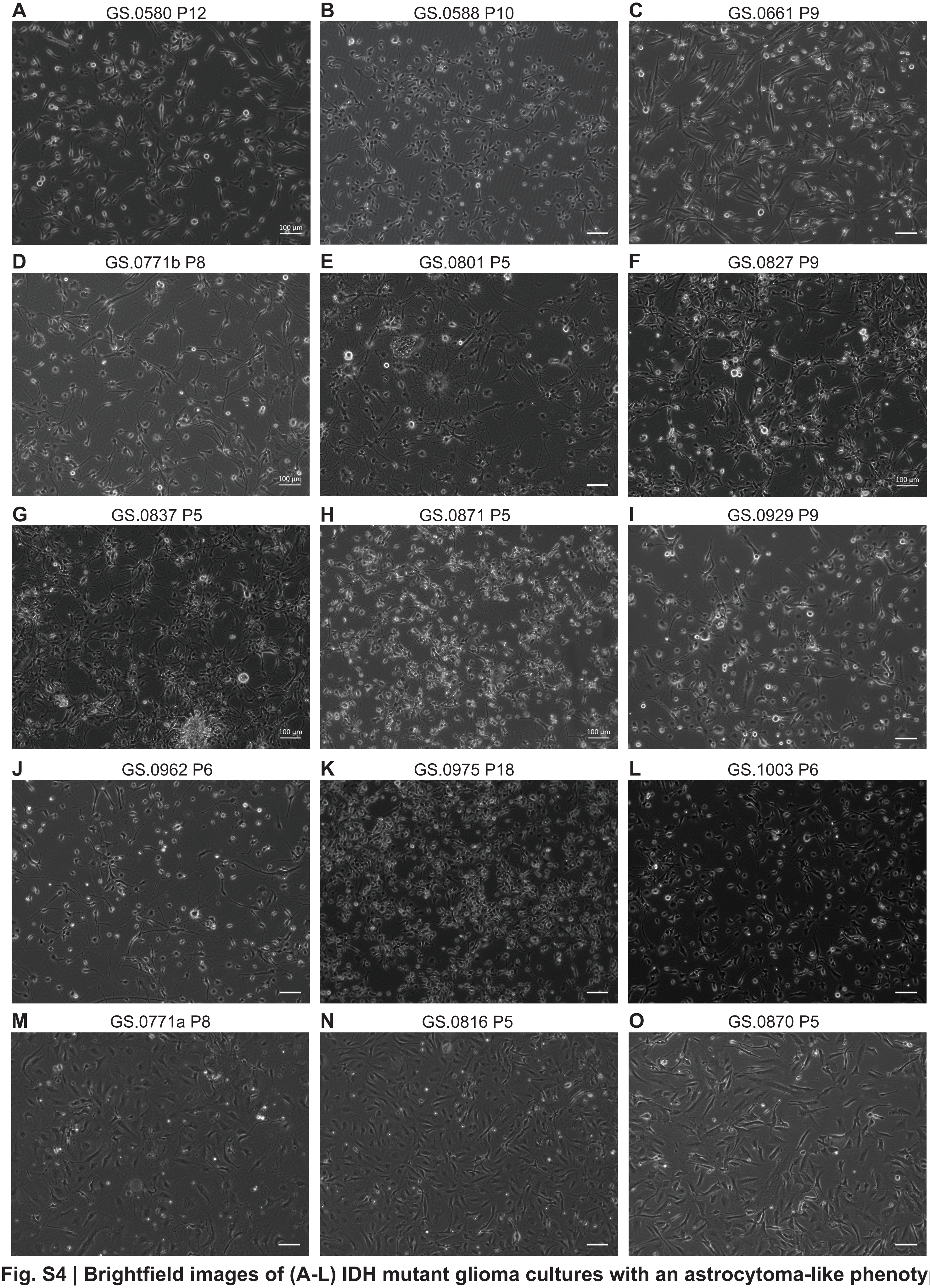


**Fig. S4 | Brightfield images of (A-L) IDH mutant glioma cultures with an astrocytoma-like phenotype and (M-O) cultures that lost the IDH mutation with a fibroblast-like phenotype**
